# Directed differentiation of human hindbrain neuroepithelial stem cells recapitulates cerebellar granule neurogenesis

**DOI:** 10.1101/2023.01.02.522503

**Authors:** Biren M. Dave, Xin Chen, Fraser McCready, Jignesh K. Tailor, James Ellis, Xi Huang, Peter B. Dirks

## Abstract

Cerebellar granule neurons (CGNs) are the most abundant neurons in the human brain and modulate cerebellar output to the motor cortex. Dysregulation of CGN development underlies movement disorders and medulloblastomas. It is suspected that these disorders arise in progenitor states of the CGN lineage, for which human models are lacking. Here, we have differentiated human hindbrain neuroepithelial stem (hbNES) cells to CGNs *in vitro* using soluble growth factors, recapitulating key progenitor states in the lineage. We show that hbNES cells are not lineage committed and retain rhombomere 1 (r1) regional identity. Upon differentiation, hbNES cells first transit through a rhombic lip (RL) progenitor state at day 7, demonstrating human specific sub-ventricular cell identities. This RL state is followed by an ATOH1^+^ CGN progenitor state at day 14. By the end of a 56-day differentiation procedure, we obtain mature neurons expressing CGN markers GABA_A_a6 and vGLUT2. These neurons generate spontaneous and evoked action potentials. A small fraction of endpoint neurons were unipolar brush cells (UBC). We noted maintenance of a RL population throughout differentiation, as is consistent with human development. We show that sonic hedgehog (SHH) promotes γ-aminobutyric acid (GABA)-ergic lineage specification and is a positive regulator of CGN progenitor proliferation. Interestingly, we observed that functional neuronal maturation is impaired by either elevated or absent SHH signaling. Impaired maturation under high SHH levels represents the potential of our system to model cerebellar tumorigenesis. Further, our data suggest a potential pro-differentiation role of SHH within a certain concentration range. Our work is, to our knowledge, the first detailed temporal characterization of the complete human CGN lineage *in vitro*. Our system recapitulates developmentally relevant progenitor states and is a new tool to model this specific cerebellar lineage, and how it may be disrupted to cause human disease.

## Introduction

Dysregulation of cerebellar granule neurogenesis underlies many human disease states, including movement disorders and medulloblastoma (Kawamura et al., 2021; Schüller et al., 2008; Vanner et al., 2014; Yang et al., 2008). Our understanding of mammalian cerebellar granule neuron (CGN) development largely depends on mouse studies, and it has long been assumed that this process is roughly equivalent in humans (Burgoyne and Cambray-Deakin, 1988). However, it has been shown that human and mouse granule lineage development differ temporally and anatomically (Aldinger et al., 2021; Haldipur et al., 2019). Specifically, the human rhombic lip (RL) persists for longer in development than its mouse counterpart and exists as ventricular zone (RL-VZ) and sub-ventricular zone (RL-SVZ) compartments. Therefore, cells underlying human disease states may not have direct equivalents in the mouse brain, preventing them from being faithfully modeled in a mouse system.

CGN differentiation begins when bone morphogenetic protein (BMP) molecules, secreted by fourth ventricle roof plate cells, act on adjacent rhombomere 1 (r1) neuroepithelium to induce RL formation (Alder et al., 1999). RL is the germinal zone of cerebellar glutamatergic neurons and is characterized by *ATOH1* and *OLIG3* expression (Akazawa et al., 1995; Lowenstein et al., 2021). *ATOH1*-expressing RL progenitors tangentially migrate over the nascent cerebellum to form a superficial external granule layer (EGL) (Ben-Arie et al., 1997; Miale and Sidman, 1961). Cells populating the EGL are called granule neuron progenitors (GNPs). Purkinje neuron-secreted sonic hedgehog (SHH) is a GNP mitogen and expands the GNP population in the EGL (Wechsler-Reya and Scott, 1999). SHH signaling protects ATOH1 from proteasomal degradation (Forget et al., 2014), allowing maintenance of the primary cilium and transduction of SHH signaling (Chang et al., 2019). Notch signaling is also active within GNPs in the EGL and prevents precocious differentiation by maintaining a proliferative state (Solecki et al., 2001). As development proceeds, promoters of *GLI1/2*, the primary downstream effectors of SHH signaling are deacetylated, repressing the SHH pathway and inducing CGN differentiation (Tiberi et al., 2014). Contemporaneously, differentiating GNPs begin to produce brain derived neurotrophic factor (BDNF) which acts in an autocrine and paracrine manner to initiate and guide radial migration to the inner granule layer (IGL) of the cerebellum (Borghesani et al., 2002). Granule neurons in the IGL extend axons up through the Purkinje layer to the molecular layer of the cerebellum. Here, their axons bifurcate into parallel fibres which form glutamatergic synapses with Purkinje neurons, which are ultimately responsible for relaying cerebellar output to the motor cortex via the deep cerebellar nuclei (Allen and Tsukahara, 1974).

Differentiation of mouse and human embryonic stem (ES) cells to cerebellar granule neurons have been reported (Behesti et al., 2021; Erceg et al., 2010; Salero and Hatten, 2007; Su et al., 2006). However, these studies focussed on characterizing end products of differentiation, rather than developmental intermediaries, from which disease states are more likely to arise (Kawamura et al., 2021; Selvadurai et al., 2020; Vanner et al., 2014). Further, they relied on co-culture of differentiating cells with mouse cerebellar cells or with growth medium conditioned by mouse cells. In such cases, it is unknown which microenvironmental factors are influencing differentiation. Functional characterization of differentiated neurons is also lacking in these works. Finally, it remains unclear if a mature neuronal state is achieved in some of these studies, due to reported expression of immature markers, such as *GLI1*, in the final differentiation timepoint (Erceg et al., 2010).

Human hindbrain neuroepithelial stem (hbNES) cells have been isolated from the embryonic hindbrain (Tailor et al., 2013). hbNES cells are long-term self-renewing stem cells that maintain their regional and temporal identities *in vitro* and have been shown to be responsive to various signaling cues. Previous studies on pluripotent stem cell derived NES cells (lt-NES) have demonstrated the developmental plasticity of these cells by differentiating them to various neuronal types *in vitro*, including motor neurons and dopaminergic neurons (Falk et al., 2012; Koch et al., 2009). However, the ability of these cells to recapitulate hindbrain developmental processes remains unknown. Given these past results, we hypothesized that hindbrain hbNES cells should possess the potential for differentiation towards the CGN lineage. If this could be achieved, a novel, genetically tractable model system could be achieved for applications in understanding human cerebellar development and pathology.

Here, we report efficient differentiation of CGNs from 4 independently derived hindbrain hbNES cell lines. Using bulk RNA sequencing (RNA-seq), single cell Multiome sequencing (scMultiome-seq), and electrophysiology, we show that differentiating hbNES cells transit through RL and GNP stages before functionally maturing. We observed recapitulation of important human specific features, such as the presence of ventricular and sub-ventricular identities during the RL stage, and the persistence of RL-like cells through to the end of differentiation. At our final day 56 differentiation timepoint, we obtain CGNs at a frequency of 86%. Additionally, we observe unipolar brush cells (UBCs) at our final differentiation timepoint. To our knowledge, this is the first report of UBCs also diverging from the granule lineage in a human context, as has been demonstrated in mice (Englund et al., 2006). We further use our system to investigate the role of SHH during granule neuron differentiation. In addition to functioning as a mitogen, high SHH levels aberrantly promote GABAergic fate. Interestingly, we uncover a potentially pro-differentiation role for SHH in the CGN lineage as spontaneous neuronal activity is diminished in its absence. Our work establishes and characterizes a human system of CGN differentiation that recapitulates key features of this process and is well suited to modeling development and diseases such as medulloblastomas and movement disorders.

## Results

### NES cells are not lineage committed, and maintain regional and temporal identity

We cultured hbNES cell lines isolated from 4 different human embryos aged gestational weeks 5 to 6 of both sexes (Tailor et al., 2013) (Supplementary table 1A). Under adherent conditions, we observed all hbNES cell lines self-organized into neural rosettes characterized by radial symmetry. hbNES cells expressed the neuroepithelial transcription factor (TF) PLZF and tight-junction protein ZO1 on their apical faces (Fig 1A). In their undifferentiated state, we confirmed that hbNES cells were not committed to the neural or glial lineage, indicated by absence of βIII-tubulin and GFAP staining, respectively (Fig 1B). Furthermore, hbNES cells maintain expression of a repertoire of neuroepithelial genes and transcription factors, such as SOX1 and SOX2 (Supplementary figs 1A, B). Together these results suggest maintenance of neuroepithelial temporal identity *in vitro*. Importantly, hbNES cells maintained a regional identity restricted to r1, based on high expression of *EN1*, *EN2* (Davis and Joyner, 1988), and *GBX2* (Joyner et al., 2000), along with low or absent expression of forebrain (*FOXG1* and *OTX1/2*) (Joyner et al., 2000; Martynoga et al., 2005) and posterior hindbrain (*HOX* genes) (Gavalas et al., 2003) markers (Fig 1C), as measured by RNA-seq. To interrogate the heterogeneity of our hbNES cell population, we profiled the transcriptional and epigenetic landscapes of 8370 undifferentiated hbNES cells in a representative line (SAI5) using scMultiome-seq. Through nearest neighbour clustering we identified 3 clusters within this population (Fig 1D). We observed ubiquitous expression of classical neuroepithelial genes *LIN28A*, *LIN28B* (Herrlinger et al., 2019), *PRTG* (Wong et al., 2010), and *SOX2* throughout these clusters (Figs 1E – H), validating our previously made observations from bulk RNA-seq and immunocytochemistry (ICC). Among this undifferentiated population, we discovered a cluster of cells where *PAX3* and *TFAP2B* were significantly differentially expressed (p = 3.25e^-115^ and 1.05e^-129^, respectively) (Figs 1I, J and Supplementary table 2C). We annotated this cluster as “Primed-NES”, based on the role of these two TFs in the differentiation of GABAergic cerebellar neurons (Zainolabidin et al., 2017). This observation corroborates a previous report that hbNES cells generate GABAergic interneurons when differentiated spontaneously (Tailor et al., 2013). To further interrogate this Primed-NES population at the epigenetic level, we performed differential TF motif activity analysis of the Primed-NES cluster relative to NES-1 and NES-2 clusters (Supplementary table 2K). In agreement with our prior observation that Primed-NES cells are inclined towards the GABAergic fate, we found that binding motifs for the GABAergic cerebellar lineage TFs TFAP2B (motif ID: MA0812.1, p = 0) and PTF1A (motif ID: MA1619.1, p = 5.13e^-148^) were significantly more active in Primed-NES cells (Fig 1M, N). However, to our surprise, Primed-NES cells also had significantly active cerebellar glutamatergic lineage TF motifs for ATOH1 (motif ID: MA1467.1, p = 5.68e^-154^) and ZIC1/ZIC2 (Aruga et al., 2002) (motif ID: 1628.1, p = 9.83e^-143^) (Fig 1K, L). Therefore, we conclude that while Primed-NES cells may be transcriptionally primed for GABAergic differentiation, epigenetically they remain plastic, and potentially can be differentiated towards the glutamatergic CGN lineage. Taken together these data demonstrate that hbNES cells retain their regional and temporal identities *in vitro*, and a subpopulation of them are primed for specific neuronal lineage differentiation.

**Figure 1.**
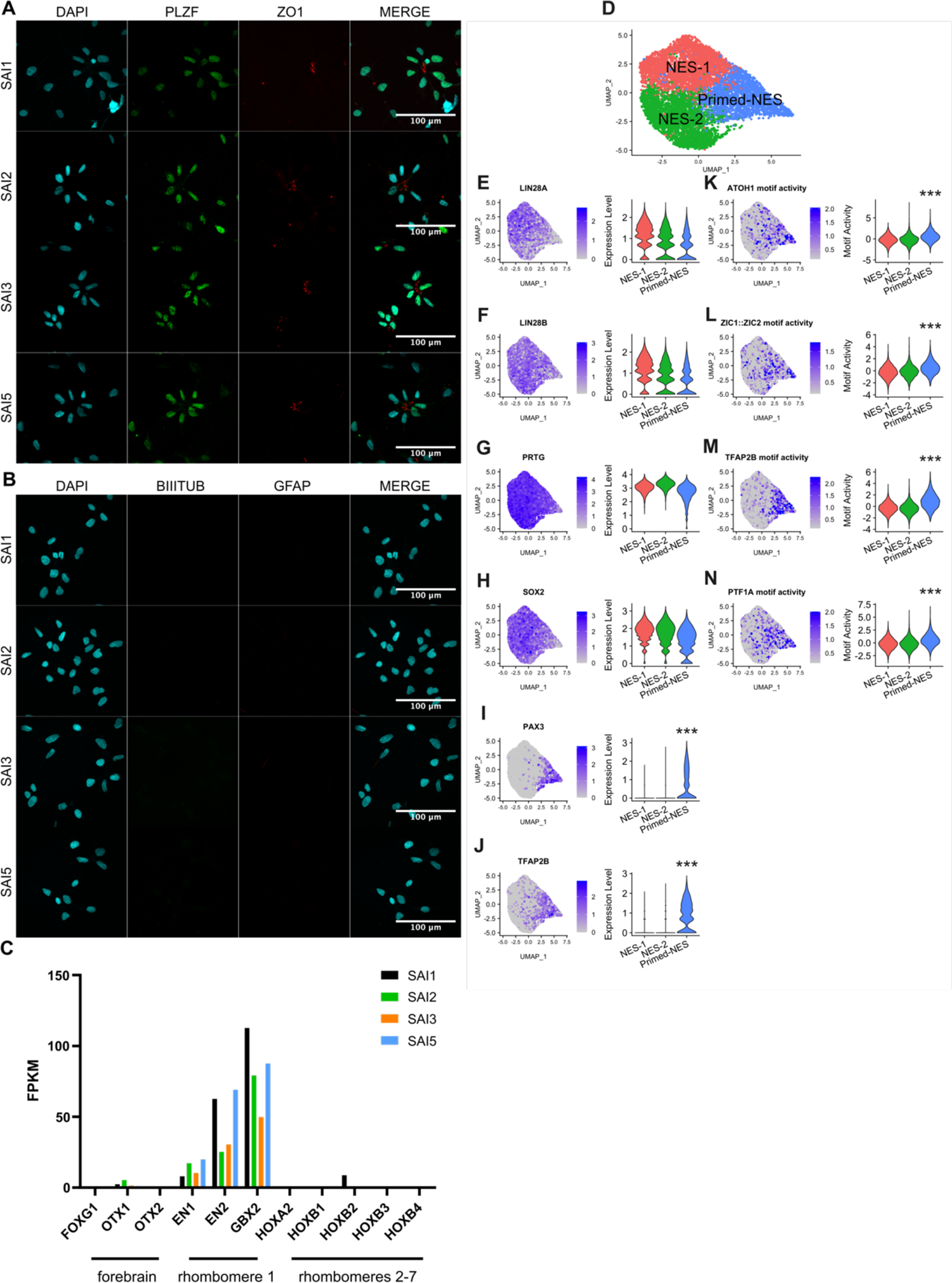
Undifferentiated NES cells stably retain regional and temporal identity *in vitro*. **A** Representative ICC images of neural rosettes formed by NES cells expressing neuroepithelial transcription factor PLZF and tight-junction protein ZO1 on their apical face. **B** Representative ICC images of neural (*β*III tubulin) and glial (GFAP) lineage marker expression by NES cells. Scale bars represent 100 μm. **C** Bulk RNA-seq expression of neural tube regional markers in NES cells. **D** Multimodal UMAP plot of undifferentiated (day 0) SAI5 NES cells. Undifferentiated NES cells express neuroepithelial genes *LIN28A* (**E**), *LIN28B* (**F**), *PRTG* (**G**), and *SOX2* (**H**). Primed-NES cells express GABAergic lineage genes *PAX3* (p = 3.25e^-115^) (**I**) and *TFAP2B* (p = 1.05e^-129^) (**J**). Transcription factor binding motifs for glutamatergic TFs ATOH1 (motif ID: MA1467.1, p = 5.68e^-154^) (**K**) and ZIC1/ZIC2 (motif ID: 1628.1, p = 9.83e^-143^) (**L**) and GABAergic TFs TFAP2B motif ID: MA0812.1, p = 0) (**M**) and PTF1A (motif ID: 1628.1, p = 9.83e^-143^) (**N**) are significantly differentially active in Primed-NES cells compared to NES-1 and NES-2 cells.

### Differentiation of independently derived hbNES cell lines to cerebellar granule neurons

Having verified the integrity of our starting hbNES cell population, we next sought to differentiate hbNES cells to CGNs to generate a model system with which to study developmentally relevant intermediate states. We modified previously reported differentiation protocols using embryonic stem (ES) cells and induced pluripotent stem (iPS) cells as their starting populations to start with hbNES cells instead (Fig 2A) (Erceg et al., 2010; Erceg et al., 2012; Salero and Hatten, 2007). First, we withdrew epidermal growth factor (EGF) and fibroblast growth factor (FGF) from the culture medium and added BMP6, BMP7, and BMP12/growth differentiation factor 7 (GDF7) for 7 days to induce a RL identity. At this timepoint, we added SHH and Jagged 1 (Jag1), a Notch2 ligand, to promote GNP formation. We also supplemented the growth medium with BDNF and Neutrophin-3 (NT3) from 7 days onward to induce neurite growth and maintain cell survival (Ghosh et al., 1994; Segal et al., 1995).

**Figure 2.**
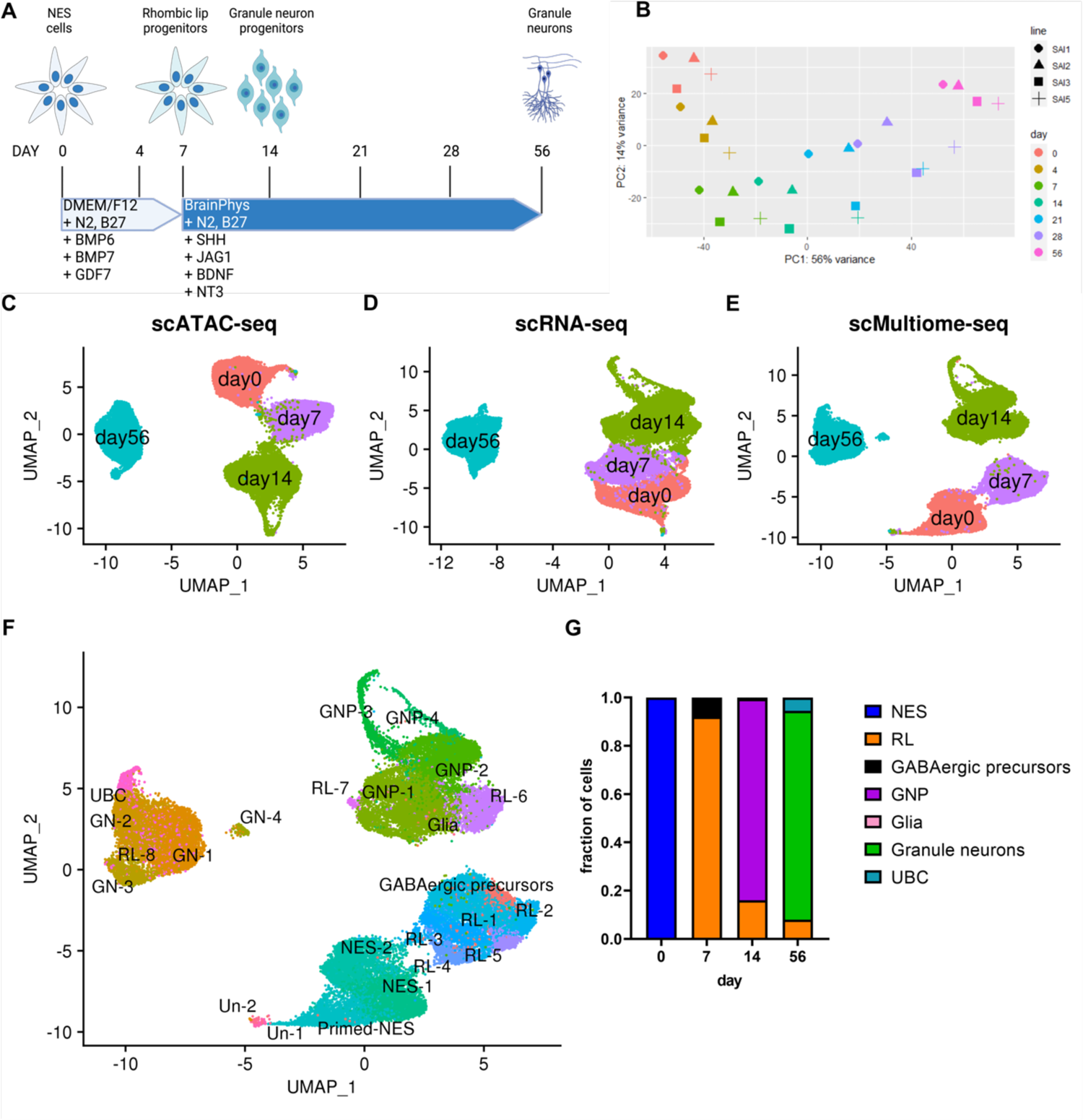
Directed differentiation of NES cells to cerebellar granule neurons. **A** Schematic of 56 day differentiation procedure. **B** Principal component analysis (PCA) plot of bulk transcriptome during differentiation. UMAP visualizations of single cell chromatin accessibilities (**C**), transcriptomes (**D**) and both chromatin accessibility and transcriptome (**E**) at days 0, 7, 14, and 56 of differentiation in SAI5 NES cells. **F** Annotation of cell clusters in figure 1E. **G** Quantification of cell types at each differentiation timepoint.

We first conducted quantitative reverse-transcriptase PCR (qRT-PCR) gene expression analysis of differentiating hbNES cells at the timepoints described in Fig 2A. We measured the expression of 24 genes spanning the entire CGN lineage from hbNES cell to mature neuron (Supplementary fig 2A). We conducted hierarchical clustering of relative expression values and noted timepoints tended to cluster together across the four different cell lines, indicating a robust response of hbNES cells to our differentiation protocol. We found no difference in the temporal sequence of differentiation between male and female hbNES cells. To further profile global transcriptomic changes at these differentiation timepoints, we conducted bulk RNA-seq and found that it corroborated the trends observed in our qRT-PCR results. Differentiation timepoints again tended to cluster together across cell lines, suggesting similar responses to differentiation in independently derived hbNES cell lines (Supplementary fig 2B, C). Two-dimensional principal component analysis (PCA) supported this conclusion and further revealed a spectrum of progenitor states that differentiating hbNES cells transit through before becoming mature neurons (Fig 2B). Further analyzing our PCA results, we noticed one set of genes, primarily contributing to principal component (PC) 1, whose expression varied monotonically with differentiation time (Fig 2B). However, genes contributing to PC2 had similar expression at days 0 and 56 but were maximally different at intermediate timepoints. We conclude from these observations that progenitor states arise and diminish during our differentiation procedure, highlighting a developmentally relevant feature of our system.

To capture the transcriptional and epigenetic changes occurring during CGN differentiation at a single cell level, we conducted scMultiome-seq at days 0, 7, 14, and 56 in a representative hbNES cell line (SAI5). This method allowed us to profile transcriptional and epigenetic landscapes from the same cell at key differentiation timepoints. We sequenced 8370, 8208, 12952 and 7914 cells passing quality control (Supplementary table 2A) at each timepoint, respectively. When we plotted chromatin accessibility (Fig 2C) or transcriptome (Fig 2D) individually, we observed a continuity of cell states through days 0, 7, and 14, with day 56 situated alone. We then combined both modalities (Fig 2E) and observed a similar trend. We noted continuity among days 0 and 7, with day 14 situated proximally and day 56 forming a distinct population. These results align with our qRT-PCR and bulk RNA-seq observations in which differentiation causes gradual changes to cell identity over time, such that day 56 cells are considerably different from those of the earlier timepoints.

To interrogate the transcriptional and epigenetic heterogeneity arising throughout differentiation, we next clustered each timepoint independently and performed differential expression (DE) and differential transcription factor (TF) binding motif activity (DA) analysis (Supplementary tables 2C – F, Supplementary fig 2C). Using markers reported in previously published datasets, we annotated each cluster (Supplementary table 2B). We then overlaid these clusters onto a multi-modal UMAP plot of all timepoints together (Fig 2F). Finally, we quantified cell types at each differentiation timepoint (Fig 2G). In our differentiation procedure, we have the emergence of a RL state at day 7, which includes human specific cell identities. This is followed by a GNP state at day 14, which matures to granule neurons at day 56 at a frequency 86%. Interestingly, we also noted the presence of UBCs, which are thought to derive from RL in mice, at day 56 at a proportion of 5%. From these data, we conclude that our differentiation procedure is robust and developmentally relevant.

### Emergence of a rhombic lip state at day 7 of granule neuron differentiation

To further characterize the RL state that arose during our differentiation procedure, we first turned to our bulk RNA-seq data. We observed that known RL markers *OLIG3* (Lowenstein et al., 2021), *ATOH1* (Ben-Arie et al., 1997), and *BARHL1* (Rachidi and Lopes, 2006) peaked in expression at day 7 (Fig 3A) and were significantly upregulated at this timepoint (Supplementary table 1F). Intrigued by this finding, we turned to our scMultiome-seq data and conducted DE analysis between all timepoints. Interestingly, three of the top differentially expressed genes at day 7 relative to other timepoints were *SLC12A2*, *TMEM132D*, and *CRABP1* (Fig 3B-D). All three of these genes have been previously reported to be expressed in the RL (Miller et al., 2014; Theisen et al., 2018; Wilson et al., 2007). Importantly, all the markers we used to characterize this timepoint as RL were either lowly or not expressed in undifferentiated hbNES cells (Fig 3). Therefore, our bulk RNA-seq and scMultiome-seq data support the emergence of a RL state at day 7 of differentiation.

**Figure 3.**
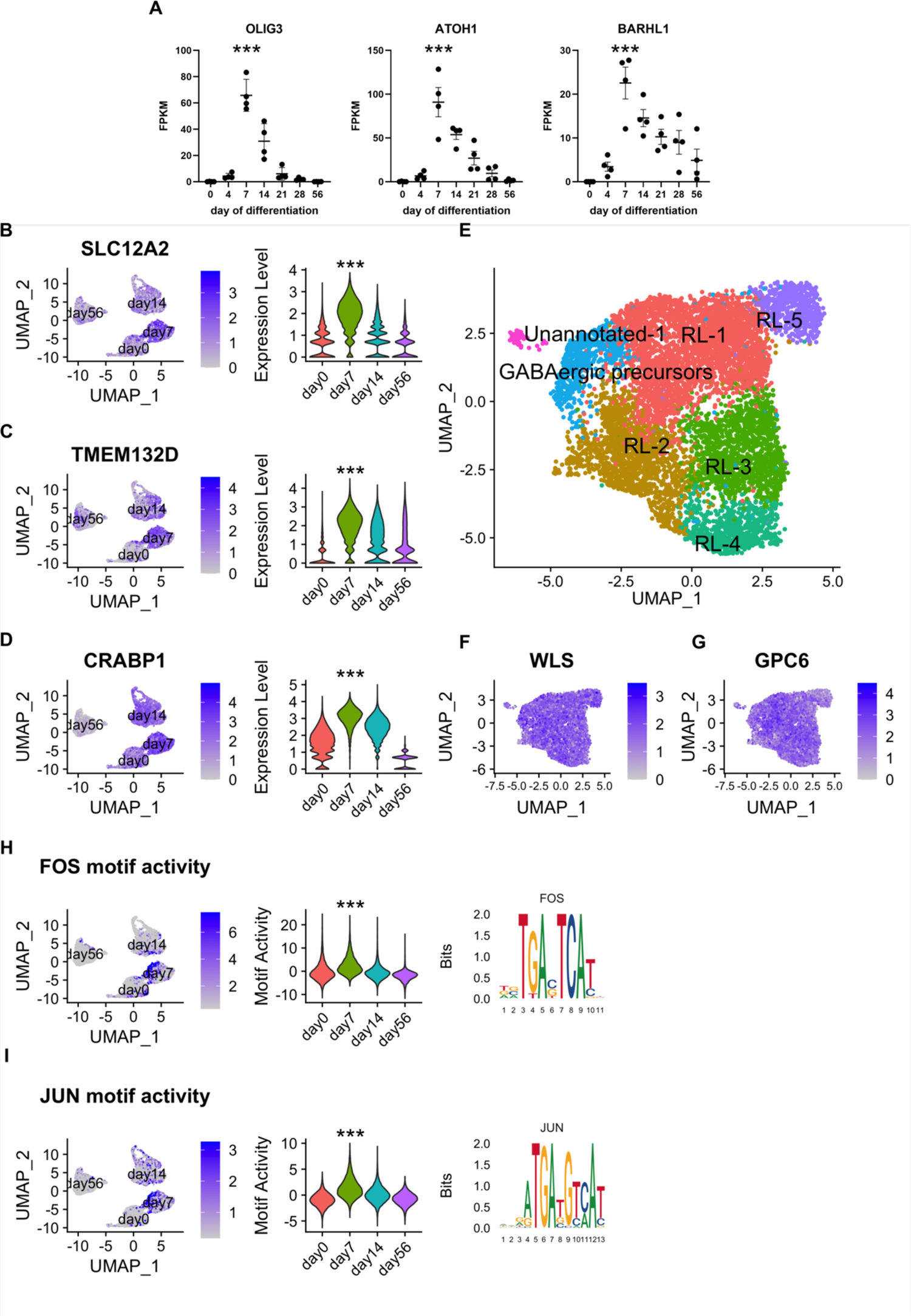
Emergence of a rhombic lip-like state at day 7 of granule neuron differentiation. **A** Rhombic lip markers *OLIG3* (log2FC = 4.95, p = 9.45e^-36^), *ATOH1* (log2FC = 3.51, p 4.29e^-13^) and *BARHL1* (log2FC = 2.60, p = 3.60e^-8^) are significantly differentially expressed at day 7 of differentiation by bulk RNA-seq compared to all other timepoints combined (see also supplementary table 1F). Among the top differentially expressed genes at day 7 by scMultiome were rhombic lip markers *SLC12A2* (log2FC = 1.63, p = 8.66e^-170^) (**B**) *TMEM132D* (log2FC = 1.22, p = 1.47e^-110^) (**C**) and *CRABP1* (log2FC = 1.56, p = 7.15e^-198^) (**D**) (see also supplementary table 2G). **E** Annotation of day 7 clusters (see also supplementary tables 2B and 2D). Expression of human rhombic lip compartment markers *WLS* (ventricular zone) (**F**) and *GPC6* (**G**) (sub-ventricular zone) at day 7. Transcription factor binding motif activities for FOS (**H**) and JUN (**I**) transcription factors.

To further characterize heterogeneity present within the day 7 population, we performed clustering, revealing 7 sub-clusters, among which we performed DE analysis to identify them (Fig 3E). Of the 7, we identified 5 as clusters of RL cells, all of which exhibited a high expression of *SLC12A2*, *TMEM132D*, and *CRABP1* (Supplementary figure 3A). We annotated a cluster of GABAergic progenitors based on *TFAP2B* and *PAX3* expression. We were left with a cluster of 88 cells which expressed no known granule lineage markers and proved challenging to assign an identity to. We designated this cluster Unannotated-1 (Un-1). However, based on differential expression of extracellular matrix genes (*DCN*, *COL6A3*, and *LUM*) and the position of this cluster adjacent to hbNES cells (Fig 2F), we suggest it represents a miniscule population of hbNES cells that are adopting a mesenchymal identity instead of differentiating.

It has recently been reported that the human RL contains a sub-ventricular compartment which is absent in mice and primates (Haldipur et al., 2019). We were interested to see if this distinction was present in our RL population, using *WLS* and *GPC6* as markers of RL-VZ and RL-SVZ, respectively (Aldinger et al., 2021). We observed ubiquitous expression of both genes in our day 7 population, suggesting presence of both VZ compartments at this differentiation timepoint.

To identify TFs that may be regulating RL identity in our system, we conducted a DA analysis of day 7 cells compared to other timepoints (Supplementary figure 2L). Interestingly, two of the most active binding motifs were those of FOS (Fig 3H) and JUN (Fig 3I). These two TFs have recently been reported to be expressed in progenitor states of the CGN lineage, and function to prevent precocious differentiation (Goodwin et al., 2021).

Therefore, our bulk RNA-seq and scMultiome-seq data suggest emergence of an RL-like state at day 7 of differentiation at approximately 92% efficiency, which includes human specific RL-SVZ identities.

### A granule neuron progenitor state follows the rhombic lip state at day 14

RL progenitor cells migrate over the surface of the cerebellum to form a superficial layer of GNPs, which proliferate in response to SHH secreted by Purkinje neurons (Miale and Sidman, 1961; Wechsler-Reya and Scott, 1999). *ATOH1* expression is maintained by GNPs until they begin to terminally differentiate (Ben-Arie et al., 1997). Given our observation of a RL-like state at day 7, we asked if it was followed by a GNP state at day 14.

We first analyzed SHH pathway activity during differentiation. *GLI1* was most highly expressed at day 14 compared to other timepoints (p = 3.18e^-11^) (Fig 4A). The maintenance of *ATOH1* expression at this timepoint (Fig 3A) suggested the presence of a GNP state, which we sought to investigate further. Since commercially available anti-ATOH1 antibodies are unreliable, we generated an ATOH1-EGFP knock-in allele in the SAI3 hbNES cell line using CRISPR-Cas9 mediated homology directed repair (Ran et al., 2013) (Supplementary Fig 4). We conducted ICC using this reporter line (Fig 4B, C), and observed that the proportion of EGFP^+^ cells was highest at day 14 (Fig 4D). Importantly, the fraction of EGFP^+^ cells that were also actively cycling (i.e., KI67^+^) was also highest at day 14. This suggests that proliferation of EGFP^+^ cells in our system is driven by SHH, which is consistent with our understanding of the effect of SHH on mouse GNPs during cerebellar development (Wechsler-Reya and Scott, 1999).

**Figure 4.**
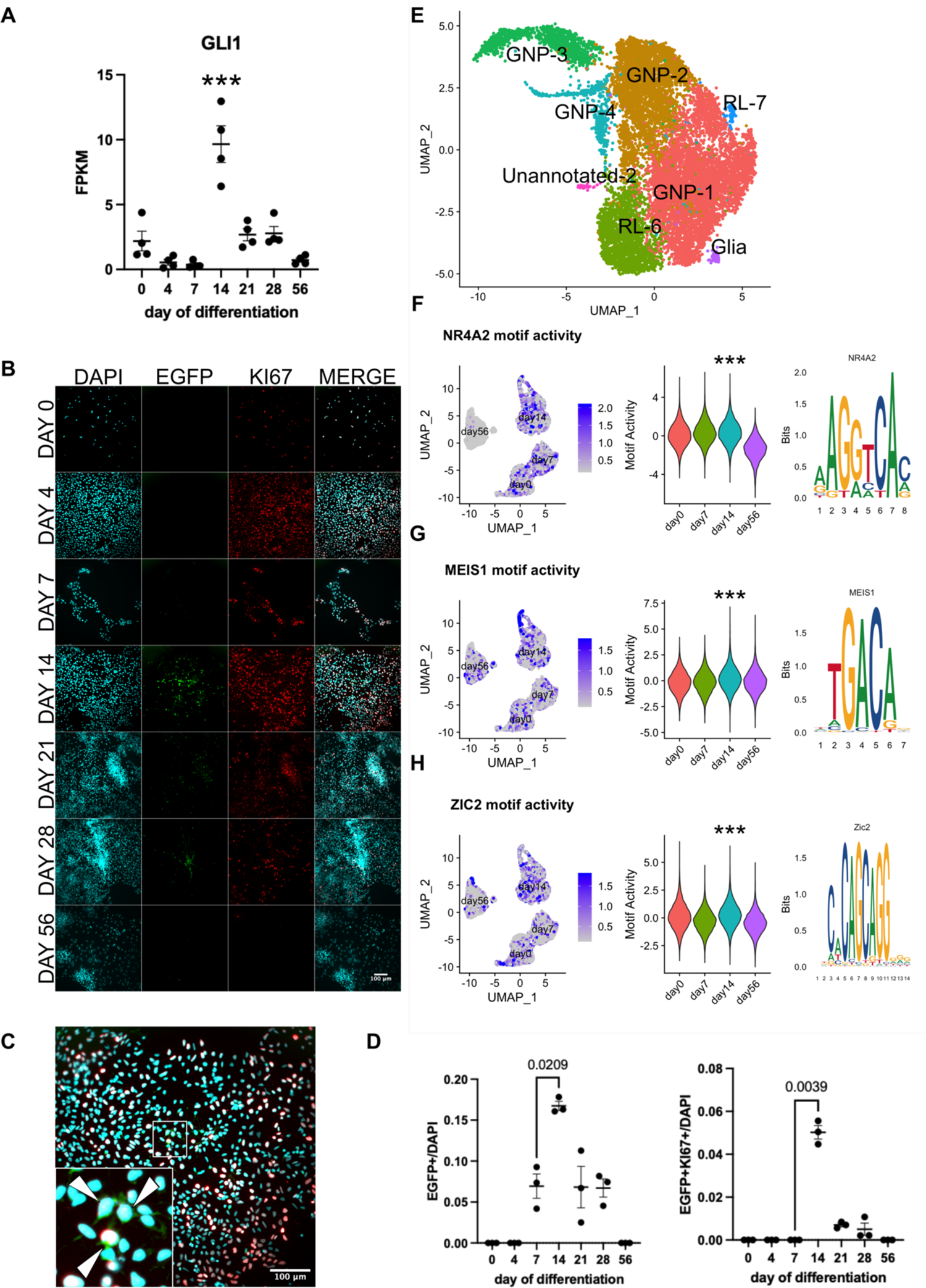
A granule neuron progenitor state follows the rhombic lip state at day 14. **A** SHH pathway effector GLI1 (log2FC = 2.84, p = 3.18e^-11^) is significantly upregulated at day 14. **B** Representative images of EGFP reporter expression during granule neuron differentiation. **C** Larger view of day 14 merged from panel B. White arrows indicate GFP^+^KI67^+^ cells. Scale bars represent 100 μm. **D** Quantification of proportion of GFP^+^ and GFP^+^/KI67^+^ cells at each differentiation timepoint (n = 3 biological replicates, t-test). **E** Annotation of day 14 cell clusters (see also supplementary tables 2B and 2E). Transcription factor binding motif activities for GNP transcription factors NR4A2 (p = 0) (**F**), MEIS1 (p = 8.01e^-150^) (**G**), and ZIC2 (p = 0) (**H**) are significantly enriched at day 14 compared to other timepoints (see also supplementary table 2M).

To dissect the cellular heterogeneity and identify active TFs within our GNP state, we turned to our scMultiome-seq data. We first performed DE analysis among timepoints, which further supported the GNP identity of day 14, based on upregulation known GNP markers *DCC*, *ZIC1*, and *ZIC3* (Aruga et al., 2002; Harter et al., 2010; Kaslin et al., 2009) (Supplementary table 2G). Interestingly, among the most differentially expressed genes at day 14 were *ID1*, *ID2*, and *ID3*: downstream effectors of BMP signaling (Ogata et al., 1993) (Supplementary table 2G, Supplementary fig 4E). This corroborates recent data showing that BMP signaling is active in human GNPs and is necessary for EGL formation in chickens (Rook et al., 2020). We next clustered day 14 independently. We identified 8 clusters, among which we performed DE analysis (Supplementary table 2E). Based on expression of known markers we identified four clusters of GNPs (based on expression of *ID1*, *ID2*, *ID3*, *MEIS2*, *ATOH1*, *DCC* and *DCX*), two RL clusters (based on *TMEM132D* and *CRABP1* expression, respectively) and one small glial cluster (based on *SCRG1* and *TSC22D* expression (Aldinger et al., 2021; Lee et al., 2022) comprising 67 cells. Like day 7, we observed a small cluster of 55 cells that proved difficult to annotate that we designated Unannotated-2 (Un-2). Interestingly, Un-2 expresses the epithelial-to-mesenchymal transition (EMT) master regulator *TWIST1*, along with other ECM genes, suggesting mesenchymal character like the Un-1 cluster.

At the epigenetic level, TF binding motifs for known GNP TFs NR4A2, MEIS1 and ZIC2 are significantly differentially active at day 14 (Fig 4F – H), lending further support to a GNP identity being present at this timepoint (Aldinger et al., 2021; Barneda-Zahonero et al., 2012; Owa et al., 2018). Together, our bulk RNAseq, scMultiome-seq and ICC data suggest RL progenitors further mature to GNPs by day 14.

### Differentiating hbNES cells mature into functional cerebellar granule neurons

As hbNES cells differentiated, we observed a gradual and concomitant decrease in the proportion of proliferating cells (Supplementary fig 5A, B). Key neuroepithelial genes were also downregulated during differentiation (Supplementary fig 5E), suggesting acquisition of a post-mitotic state and loss of stemness, respectively. Contemporaneously, differentiating cells started to extend βIII-tubulin^+^ neuronal arbors, which increased in complexity with differentiation time (Supplementary fig 5C). Importantly, we observed few or no GFAP^+^ cells during all timepoints, suggesting that our differentiation is almost entirely neuronal. We additionally stained for NEUN, a marker of post-mitotic neurons. We first detected NEUN expression at 28 days, and it remained present at 56 days further suggesting attainment of a mature neuronal state. Furthermore, we detected PAX6, a known marker of the CGN lineage was present throughout differentiation (Swanson et al., 2005) (Supplementary fig 5D).

**Figure 5.**
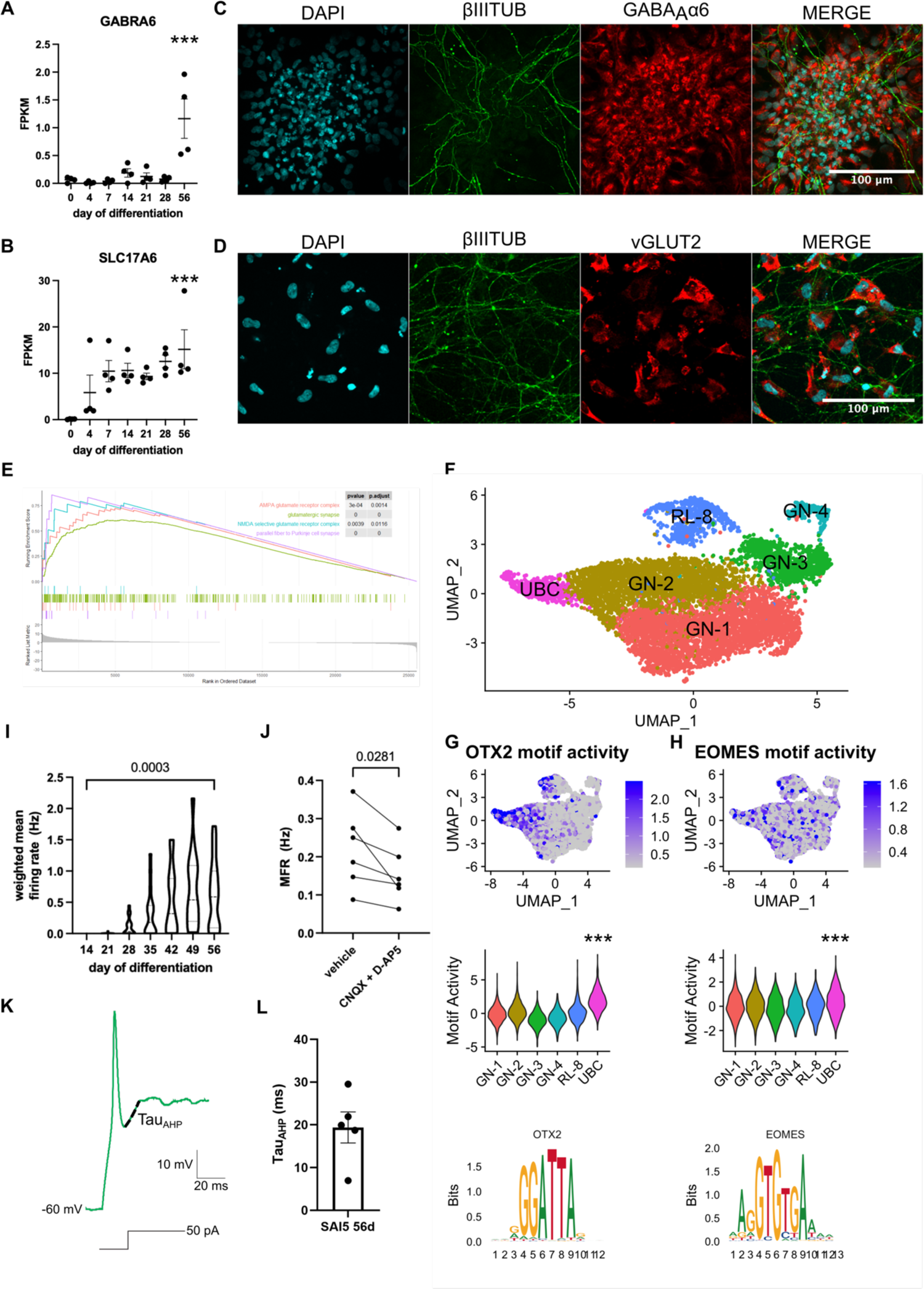
NES cell differentiation culminates in functional granule neurons. Bulk RNA-seq expression of mature granule neuron markers (**A**) *GABRA6* (log2FC = 4.82, p = 8.32e^-4^) and (**B**) *SLC17A6* (log2FC = 7.00, p = 1.81e^-41^) is significantly upregulated at day 56 of differentiation relative to day 0 (see also supplementary table 1J). Representative ICC images of GABA_A_*α*6 (**C**) and vGLUT2 (**D**) at day 56 of differentiation in the SAI5 NES cell line (scale bars represent 100 μm). (**E**) Significantly upregulated pathways identified through GSEA analysis of differentially expressed genes between days 56 and 0. (**F**) Annotation of day 56 cell clusters (see also supplementary table 2B and 2F). Activity of (**G**) OTX2 (p = 0) and (**H**) EOMES (p = 3.10e^-10^) motifs among day 56 clusters. Quantification of weighted mean firing rate (wMFR) (**I**) during differentiation as measured by microelectrode array (MEA) (n = 16 biological replicates, t-test). See Supplementary figure 5F for representative raster plots of neural activity. Quantification of mean firing rate (MFR) (**J**) upon vehicle or CNQX and D-AP5 treatment (n = 6 biological replicates, t-test). **K** Representative trace of an evoked action potential in day 56 SAI5 neurons. Dashed line indicates recovery from after-hyperpolarization (Tau_AHP_), fitted with a single exponential decay function. **L** Quantification of Tau_AHP_ in day 56 SAI5 neurons (n = 5).

We next sought to determine if the cells we had obtained at our day 56 endpoint were indeed CGNs. Using our bulk RNA-seq data, we conducted DE analysis between days 0 and 56. We noticed two genes expressed by mature CGNs, *SLC17A6* and *GABRA6* to be significantly upregulated at day 56 (Fig 5A, B) and we further confirmed expression of these markers at the protein level by ICC (Fig 5C, D). We next performed gene set enrichment analysis (GSEA) of the significantly DE genes between these two timepoints (Fig 5E, supplementary table 1J – M). We noticed several pathways pertaining to glutamatergic synaptic transmission were significantly enriched at day 56 relative to day 0. Notably, one of these terms was “parallel fiber to Purkinje cell synapse,” which refers to the glutamatergic synapse formed by CGN axons onto Purkinje neuron dendrites in the cerebellar molecular layer. These results suggest that the cells we obtain at the end of our differentiation have a mature CGN identity.

We further characterized the day 56 population at a single cell level. Relative to other timepoints, CGN markers *RELN*, *GRIK2*, and *NFIA* (Aldinger et al., 2021; Goldowitz et al., 1997) were significantly upregulated at day 56 (Supplementary table 2G), providing further evidence of a CGN identity. We then clustered day 56 independently and discovered six clusters, among which we performed differential expression analysis (Supplementary table 2F). We were able to annotate these clusters based on the expression of known markers as either CGNs (based on expression of *ROBO2*, *RELN*, *GRM8*, *NFIB*, *FSTL4* (Aldinger et al., 2021), UBCs (*TRPM3*, *OTX2* (Vladoiu et al., 2019), and RL (*TOP2A*, *ASPM*, *CENPE* (Aldinger et al., 2021) (Fig 5F, Supplementary table 2B). The overall output of our differentiation procedure was 86% granule neurons, 8% RL, and 6% UBCs (Fig 2G). We corroborated the presence of UBCs at this timepoint by conducting differential TF motif activity analysis among day 56 clusters. We found significantly enriched motif activity for OTX2 and EOMES in the UBC cell cluster (Fig 5G, H). This observation confirms reports from mouse development that UBCs arise from the granule lineage (Englund et al., 2006), and to our knowledge is the first confirmation of this developmental feature in a human context. The persistence of a RL population at day 56 confirms latest reports of human cerebellar development, in which the RL is a long-lived germinal zone, present for up to the first two years of life (Haldipur et al., 2019).

To assess the functional potential of our model, we performed electrophysiological measurements of neuronal activity. We differentiated hbNES cells on microelectrode arrays (MEA) to record spontaneous extracellular activity at a population level every week from days 14 to 56. Extracellular spikes were first detected at day 28 and continued to be recorded with increasing frequency through day 56 (Fig 5I), representing functional/neuronal activity. To determine whether glutamate was the neurotransmitter responsible for driving the spontaneous activity of our neurons, we pharmacologically blocked all post-synaptic glutamate receptors with small molecules CNQX (20 μM) and D-AP5 (100 μM), causing a significant decrease in mean firing rate (Fig 5I). We next assessed neuronal function at a single cell level using current-clamp whole cell electrophysiology. We observed that neurons at day 56 can generate single evoked action potentials (Fig 5K), however they remain depolarized after stimulation and do not fire repeatedly. This can be attributed to a combination of lack of supporting cell types, absence of neural circuit integration (Mattugini et al., 2019), and absence of appropriate voltage-gated potassium ion channels required to sufficiently hyperpolarize the cell membrane. We quantified recovery from after-hyperpolarization (Tau_AHP_) on neurons we patched and observed that they were in agreement with Tau_AHP_ values previously described for human ES cell derived neurons (Schrenk-Siemens et al., 2015). Taken together, our data describe that the predominant output of our differentiation protocol is CGNs that are capable of spontaneous and evoked neural activity *in vitro*.

### SHH functions as a mitogen and promotes GABAergic identity within the cerebellar granule neuron lineage

Having generated and characterized a system of CGN development, we were next interested in studying the role of SHH in this process. Previous reports have shown that SHH acts as a mitogen for GNPs (Wechsler-Reya and Scott, 1999) and is also required for cerebellar GABAergic neural development in mice (Huang et al., 2010). Therefore, we hypothesized that SHH plays a similar role in human CGN differentiation. We designed an experimental approach in which we exposed differentiating hbNES cells to 3 different concentrations of SHH in the growth medium starting at day 7 (Fig 6A). We noticed a dose-responsive relationship between SHH target *GLI1* expression and SHH concentration, verifying our experimental approach (Fig 6B). Interestingly, we observed that SHH does not appear to be required for commitment to the granule lineage, as *ATOH1* expression was unchanged among all three SHH conditions (Fig 6C). This result reinforces the idea that granule lineage commitment likely occurs at a RL stage in a human context before progenitor cells migrate to form the EGL. Further, we noted a dose-responsive relationship between *SOX2* expression and SHH concentration (Fig 6D). Previous work demonstrating that GLI transcription factors regulate *SOX2* expression provides a possible mechanism for this observation (Pietrobono et al., 2021), and is consistent with our previous identification of a SOX2^+^ population persisting in the mouse EGL (Selvadurai et al., 2020). Next, we sought to investigate the role of SHH as a mitogen during human CGN differentiation. Using our ATOH1-EGFP reporter line, we performed KI67 staining at day 14 of differentiation to quantify the fraction of proliferating cells. We observed that the fraction of GFP^+^ cells was similar under all 3 concentrations of SHH, (Supplementary fig 6A, C), confirming our qRT-PCR result. On day 14, we observed a positive correlation between concentration of SHH in the growth medium and proportion of GFP^+^ cells that were also KI67^+^. This result further confirms the physiological relevance of our system, as it demonstrates that SHH acts as a mitogen of GNPs.

**Figure 6.**
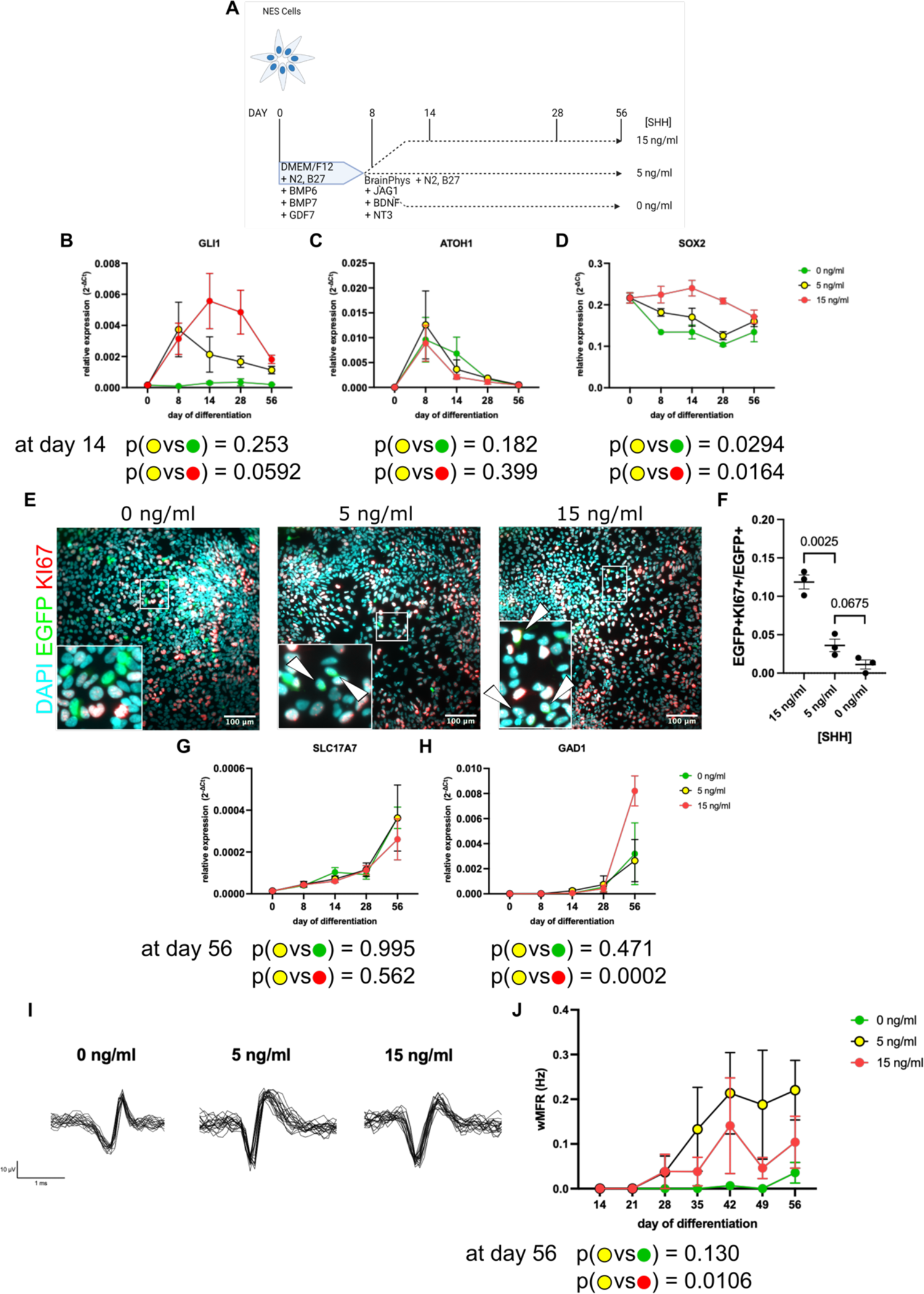
SHH functions as a GNP mitogen and encourages GABAergic lineage specification. **A** Experimental approach to test the effect of SHH on granule neuron differentiation. qRT-PCR expression of (**B**) *GLI1*, (**C**) *ATOH1*, and (**D**) *SOX2* in differentiating NES cells in varying concentrations of SHH (n = 3 biological replicates, t-test). **E** Representative ICC images of GFP and KI67 expression at day 14 of differentiation under various concentrations of SHH. White arrows indicate ATOH1^+^KI67^+^ cells in image insets. **F** Quantification of GFP^+^KI67^+^ cell proportion (n = 3 biological replicates, t-test). **G** Representative waveforms spontaneously generated by neurons differentiated under all three SHH concentrations (n = 20 for each condition). **H** Quantification of weighted mean firing rate (wMFR) in differentiating NES cells by MEA. Expression of (**I**) glutamatergic (*SLC17A7*) and (**J**) GABAergic (*GAD1*) markers during differentiation by measured by qRT-PCR (n = 3 biological replicates, t-test). All scale bars represent 100 μm.

To test if SHH promotes the GABAergic fate in the CGN lineage, we analyzed the expression of *SLC17A7* and *GAD1*, glutamatergic and GABAergic markers respectively during differentiation using qRT-PCR (Figs 6G, H). Neurons derived under all 3 conditions showed similar extents of arborization (Supplementary fig 6D). However, we observed that the GABAergic marker *GAD1* was significantly upregulated in neurons differentiated in 15 ng/ml of SHH compared to 5 ng/ml (Fig 6H). *SLC17A7* expression was also decreased in neurons differentiated under high SHH, though not significantly (Fig 6G). We then asked if these fate differences manifest functionally. Using MEA methodology, we observed that although functional neurons were generated under all conditions (Fig 6I), those exposed to moderate levels of SHH had the highest levels of spontaneous activity (Fig 6J). Therefore, we propose a model whereby high SHH activity potentiates an immature cell state and the GABAergic lineage, leading to less spontaneous neuronal activity at a population level. However, in the absence of SHH, functional neuronal maturation is also impaired perhaps suggesting a pro-differentiation role of SHH within a specific concentration range.

## Discussion

### Differentiation of human hbNES cells recapitulates developmentally relevant progenitor states in the cerebellar granule lineage with high efficiency

Diseases of the granule lineage arise in progenitor states for which human models have hitherto been lacking. Through the directed differentiation of independently derived human hindbrain hbNES cells, we present a method to efficiently obtain RL and GNP cell states. Extended differentiation of hbNES cells produces functional mature CGNs at 56 days. Importantly, we observed recapitulation of a human specific RL-SVZ compartment, and persistence of a RL identity to day 56. These findings are consistent with recent reports of maintenance of a RL progenitor pool for as long as the first two years of life (Aldinger et al., 2021; Haldipur et al., 2019).

Though the CGN lineage was the predominant output of our differentiation, we observed small populations of GABAergic progenitors and glia at days 7 and 14 respectively. The likeliest explanation for these populations is that a minority of very strongly GABAergically primed hbNES cells commit to the cerebellar ventricular zone lineage upon EGF and FGF withdrawal. We propose that such cells first generate GABAergic VZ progenitors at day 7, and then glia at day 14, which are known to derive from the VZ. We also observed small clusters of cells at days 7 and 14 which proved difficult to annotate but expressed markers suggestive of mesenchymal identity. Given the positions of these clusters adjacent to undifferentiated hbNES cells on our global UMAP (Fig 2F), we speculate that they may represent small populations of hbNES cells that were resistant to differentiation and instead adopted mesenchymal character.

Despite the absence of a 3-dimensional environment and lack of interaction with other supporting cell types which facilitate CGN differentiation *in vivo* (Consalez et al., 2021; Leto et al., 2016), we nonetheless recapitulate the full temporal sequence of human CGN development. By providing an in-depth resource of human CGN development at single-cell resolution, our work complements recent studies of human cerebellar development using organoids and primary tissue (Aldinger et al., 2021; Nayler et al., 2021). We present a new method by which hypotheses pertaining to CGN development and disease can be functionally tested.

### Human granule neuron differentiation is a powerful model to study development and disease

The ability to isolate and study the contributions of individual growth factors is important to understanding developmental processes. Such experiments are technically challenging to perform *in vivo*, because it may not be feasible to specifically modulate the level of growth factor a certain cell type is exposed to in a whole organism. Additionally, individual growth factors may regulate several developmental processes simultaneously, which may confound the process of interest. Therefore, *in vitro* systems such as ours are best suited to answer such questions. We leveraged this strength of our system to test the role of SHH during CGN differentiation, revealing an intriguing role of SHH in lineage determination and functional maturation. Our observation that elevated SHH signaling potentiates an immature cell state and inhibits functional neuronal maturation is consistent with the finding that constitutive SHH pathway activation can transform granule lineage cells to seed medulloblastoma (Schüller et al., 2008; Selvadurai et al., 2020; Vanner et al., 2014; Yang et al., 2008).

The ability of embryonically derived hbNES cells to stably maintain their regional and temporal identities *in vitro* represents perhaps the most powerful aspect of this system. Since all cerebellar tissue arises from r1, it is conceivable that with application of the appropriate combination of growth factors for an optimal amount of time, all human cerebellar lineages can be derived *in vitro*, thereby generating experimental models for the full spectrum of cerebellar pathologies.

An exciting observation of our day 56 population was the emergence of UBC-like cells. UBCs are glutamatergic interneurons found in the cerebellar granule layer. To our knowledge, our work is the first derivation of this neuronal type in a human context. In mice, UBCs derive from the RL, albeit at later developmental timepoints than CGNs. Here, we demonstrate a similar human developmental origin for human UBCs and lay the groundwork for further studying this important cell type in a human context. Considering recent reports implicating UBC lineage stem and progenitor cells as origins for group 3 and group 4 medulloblastoma (Hendrikse et al., 2022; Smith et al., 2022), further study of this lineage and its use in tumour modeling will be of great clinical interest.

Overall, we define a complete temporal sequence of human CGN lineage development *in vitro*, highlighting the cell state transitions from a primed stem cell population to committed neural progenitors, to differentiated and functional granule neurons. By defining the distinct steps of lineage transition, this system provides an opportunity to model the timing and cellular context of disease associated alterations and their effects on human cerebellar cellular maturation.

## Materials and methods

### NES cell culture

hbNES cells isolated from primary human fetal sources aged gestational weeks 5 to 6 (Carnegie stages 15 – 17) (Tailor et al., 2013) were grown in DMEM/F12 (Wisent, cat# 319-075-CL), 1X N2 (homemade), 0.01X B27 (Life Technologies, cat# 17204-44), 10 ng/ml rhEGF (Sigma, cat# E9644) and 10 ng/ml bFGF (STEMCELL Technologies, cat# 02634) in a humidified 37°C, 5% CO2 incubator on Corning Primeria or Falcon tissue culture vessels coated with Poly-L-Ornithine (Sigma, cat# P4957-50ML) and Laminin (Sigma, cat# L2020-1MG). hbNES cell lines tested negative for Mycoplasma. When cells neared confluency, they were dissociated with TrypLE express (Life Technologies, cat# LS12605028) for 5 min. The dissociation was quenched with DMEM/F12, cells were pelleted by centrifugation at 400 rcf for 5 min and then replated at desired density. Cells were cryopreserved in growth medium supplemented with 10% DMSO at −80°C or in liquid nitrogen. Growth medium was partially replaced every 24 hours.

### ATOH1-EGFP knock-in reporter line generation

A guideRNA sequence directing Cas9 cleavage 2 bp away from the desired site of knock-in was found using Benchling (5’ – TGACTCGGATGAGGCAAGTT – 3’, on target score: 25.8, off target score: 75.3). Sense (CACCGTGACTCGGATGAGGCAAGTT) and anti-sense (AAACAACTTGCCTCATCCGAGTCAC) oligonucleotides were hybridized and ligated into pSpCas9(BB)-2A-GFP (PX458) (addgene plasmid# 48138; gift from Feng Zhang) as previously described (Ran et al., 2013). A homology repair template was designed and ordered from VectorBuilder. hbNES cells were nucleofected using the Amaxa 4D-Nucleofector X-unit system (Lonza Bioscience, cat# AAF-1002X) when they had grown to 75% confluency. 2 x 10^6^ cells were pelleted and resuspended in 100 µL of P3 solution containing 2.5 µg of PX458 and 2.5 µg of homology template. The cell suspension was transferred into a 100 µL nucleofection cuvette (Lonza Bioscience, cat# V4XP-3024) and nucleofected with code DN100, which has previously been demonstrated to deliver plasmid cargo with high efficiency to adherent neural stem cells (Bressan et al., 2017). After nucleofection, cells were transferred to one well of a six well plate and allowed to recover for six days. Cells that had undergone homologous recombination were selected for with 1 mg/ml G418 (Invitrogen cat# 11811023). Presence of the knock-in allele was detected by PCR (see primer sequences below). Homologous recombinants were further isolated by twice sorting the population for mCherry^+^/DAPI^-^ cells on a Sony MA900 cell sorter.

#### Genotyping PCR primers

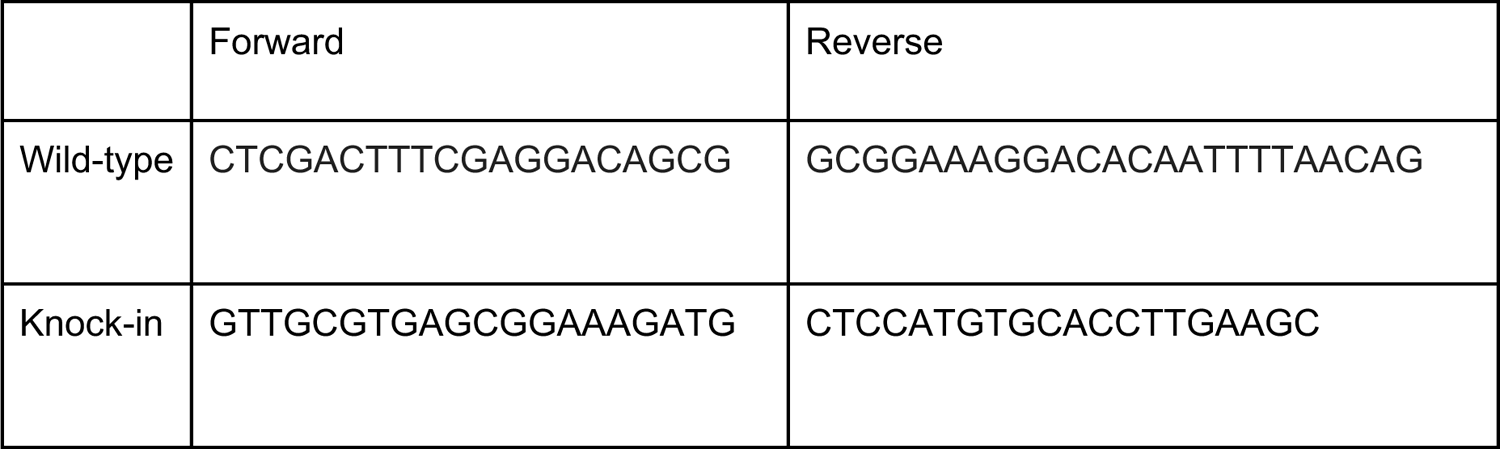

### Granule neuron differentiation

6 x 10^5^ hbNES cells were plated in growth medium on Poly-L-Ornithine/Laminin coated 6 cm plates on the day prior to differentiation, and 2.5 x 10^4^ hbNES cells were plated on coated glass coverslips. On day 0 of differentiation, growth medium was completely aspirated and replaced with Step 1 differentiation medium: DMEM/F12, 1X N2, 0.01X B27, 2.5 ng/ml BMP6 (R&D Systems, cat# 507-BP-020/CF), 5 ng/ml BMP7 (R&D Systems, cat# 354-BP-500/CF), and 50 ng/ml BMP12/GDF7 (R&D Systems, cat# 8386-G7-050). On day 4, differentiating cells were fully confluent and split 1:3 into Step 1 differentiation medium. On day 7, differentiating cells were confluent again and split 1:3-1:4 into Step 2 differentiation medium: BrainPhys Neuronal Medium (STEMCELL Technologies, cat# 05790), 1X N2, 0.01X B27, 2.5 ng/ml BMP6, 5 ng/ml BMP7, 50 ng/ml BMP12/GDF7, 5 ng/ml SHH (R&D Systems, cat# 8908-SH), 2 ng/ml JAG1 (1277-JG-050, cat# 1277-JG-050), 10 ng/ml NT3 (R&D Systems, cat# 267-N3-025/CF) and 100 ng/ml BDNF (R&D Systems, cat# 248-BDB-01M/CF). From days 0 to 28, medium was partially replaced every 48h. After day 28, medium was partially replaced twice per week. For the SHH dependence experiment two additional formulations of step 2 differentiation medium were prepared with 15 ng/ml SHH or 0 ng/ml SHH.

### Quantitative real-time PCR (qRT-PCR)

Total RNA was extracted from cell pellets using the Qiagen RNeasy Plus Mini Kit (Qiagen, cat# 74134). cDNA was synthesized with Transcriptor Reverse Transcriptase (Roche, cat# 3531287001), using an oligo(dT)_18_sequence. Reactions were carried out in either 96 well (Bio-rad, cat# HSP9661) or 384 well (Thermo Scientific, cat# 43-098-49) format using SsoFast EvaGreen Supermix (Bio-rad, cat# 1725202) or PowerUp SYBR Green Master Mix (Thermo Scientific, cat# A25778), respectively. Primer specificity and amplification efficiency for genes of interest were verified by performing reactions on a dilution series of human reference cDNA (Takara, cat# 636693). Gene expression relative to GAPDH and RPLP0 house-keeping genes was calculated in Microsoft Excel, and data were visualized using GraphPad Prism and Morpheus.

### Immunocytochemistry

Cells grown on glass coverslips were fixed in 10% formalin for 10 minutes at room temperature. Cells were blocked and permeabilized in a PBS solution of 0.1% Triton X100 and 5% normal goat serum (NGS) for 1 hour. Cells were incubated with primary antibody at the indicated concentration for either 1 hour at room temperature or overnight at 4°C, washed trice, and then incubated with secondary antibody for 1 hour at RT. Coverslips were mounted on glass slides with fluorescence mounting medium. (Dako, cat# 34538). Cells were imaged on a Zeiss Epifluorescence or Leica SP8 Lightning STED confocal microscope at the SickKids Imaging Facility. Images were analyzed using ImageJ and quantified in GraphPad Prism.

#### Primary antibodies

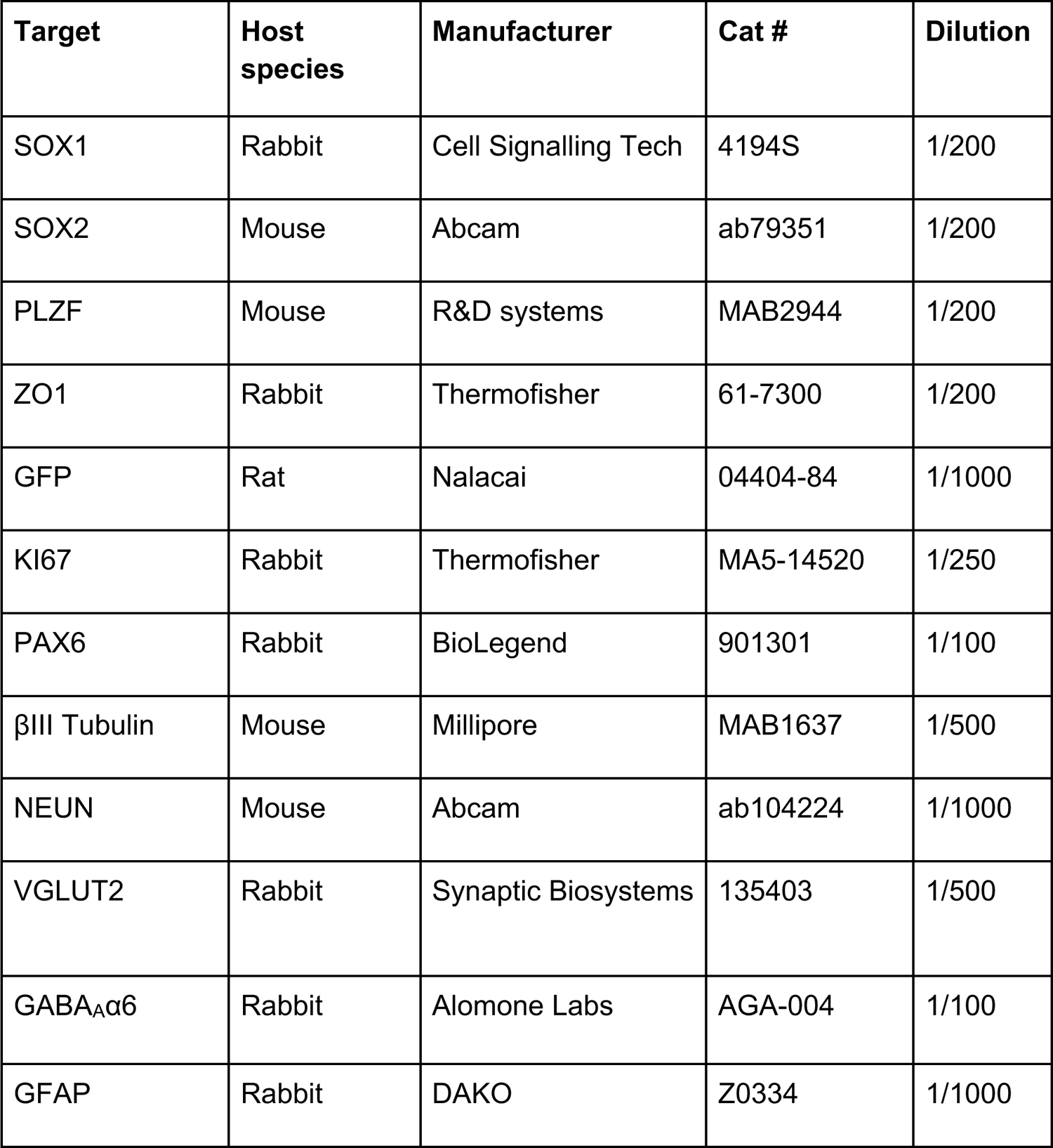

#### Secondary antibodies

| Target | Host species | Fluorophore | Manufacturer | Cat # |
| --- | --- | --- | --- | --- |
| Mouse IgG | Goat | 488 | Invitrogen | A11029 |
| Rabbit IgG | Goat | 647 | Invitrogen | A21244 |
| Rat IgG | Goat | 488 | Invitrogen | A11006 |

### Microelectrode array (MEA)

24 or 48 well CytoView plates with 16 recording channels per well were obtained from Axion Biosystems (cat #s M384-tMEA-24W, M768-tMEA-48W). MEA plates were coated with Poly-L-Ornithine overnight, washed four times with water, and then coated with laminin for at least 24 hours prior to cell plating. On day 7 of differentiation 1.5 x 10^4^ cells were plated into each well in a 70 µL droplet, containing laminin in a 1:100 dilution. Cells were allowed 60 min to adhere after which medium was topped up to 200 µL (48 well plate) or 500 µL (24 well plate). MEA plates were recorded for spontaneous activity every 7 days from day 14 of differentiation to day 56 using the AxIS software (version 2.5.2.1) on the Maestro Original system (Axion Biosystems). Recordings were performed at a constant temperature of 37°C, and plates were allowed to rest undisturbed in the pre-warmed reader before spontaneous neural activity was recorded for an additional 5 minutes. Pharmacology experiments were performed at day 56. CNQX (Biotechne, cat# 0190) and D-AP5 (Biotechne, cat# 0106) were reconstituted in DMSO and artificial cerebrospinal fluid (ACSF), respectively. Cells were first equilibrated in growth medium containing vehicle (i.e., DMSO and D-AP5) for 10 minutes before activity was recorded for 5 minutes. Medium was then completely removed and cells were washed once with PBS before being equilibrated in growth medium containing CNQX (20 μM) and D-AP5 (100 μM) for 10 minutes, before activity was recorded for 5 minutes. Recording channels were sampled at a frequency of 12.5 kHz and bandpass filtered from 200 Hz to 3 kHz. Spikes were then detected using an adaptive threshold-crossing method, with the threshold set to 6 times the standard deviation of the estimated noise of each channel. Offline analysis was conducted using the Neural Metric Tool (Axion Biosystems, version 3.1.7). Weighted mean firing rate (wMFR) was calculated as the mean firing rate normalized by the number of active electrodes in each well. Electrodes were considered active if at least 5 spikes per minute were detected. Single channel burst detection was performed using the Poisson surprise algorithm, with the “minimum surprise” parameter set to 3. Spike and burst metrics obtained using the Neural Metric Tool were subsequently visualized using GraphPad Prism. Extracellular action potential waveforms were obtained from AxIS software and visualized in GraphPad Prism.

### Whole cell patch-clamp electrophysiology

NES cells differentiated to day 56 on glass coverslips were transferred to a recording chamber filled with bath solution. The bath solution consisted of (in mM) 140 NaCl, 5 KCl, 2 CaCl_2_, 2 MgCl_2_, 10 Glucose, and 10 HEPES (pH adjusted to 7.4 with NaOH). Patch pipettes for recording, with resistance of 3-4 MΩ, were filled with intracellular solution consisting of (in mM) 123 K-Gluconate, 10 KCl, 1 MgCl_2_, 1 EGTA, 10 HEPES, 0.1 CaCl_2_, 1.2 ATP, and 4 Glucose (pH adjusted to 7.2 with KOH). Whole-cell recordings were made with a MultiClamp 700B amplifier paired with a Digidata 1440A digitizer and acquired online using the pClamp10 software package. In current-clamp experiments, cells were injected with minimal current to maintain their voltage at around −60mV and current steps were injected starting with −50 pA for 500 ms, increasing each sweep by 50 pA to a final amount of 650 pA. Representative trace in the figure demonstrates that the action potential (AP) was elicited at 50 pA current injection. All experiments were performed at room temperature. AP traces were quantified and graphed using GraphPad Prism and Adobe Illustrator.

### Bulk RNA-sequencing and analysis

250 ng of RNA from each sample were used for library preparation using the NEBNext Ultra II Directional poly(A) mRNA library prep kit (cat #E7760S). Paired-end sequencing (2 x 100 bp) was then conducted on an Illumina Novaseq S1 flowcell. Reads were mapped to release 36 (GRCh38.p13) of a human reference transcriptome using Salmon (Patro et al., 2017). Downstream analysis was performed in R using tximport (Soneson et al., 2016) and transcript abundances were log-normalized prior to differential expression analysis with the DESeq2 package (Love et al., 2014). For DE analysis, a log fold change of 1 and a false discovery rate of 0.05 were set. P-values were calculated using the Wald test and were adjusted for multiple hypothesis testing. Gene set enrichment analysis was performed using the ClusterProfiler package in R (Wu et al., 2021).

### Single-cell multiome sequencing and analysis

Cryopreserved cell suspensions were prepared in growth medium supplemented with 10% DMSO. Cells were thawed and nuclei were isolated according to protocol CG000365, Revision C by 10X Genomics (Nuclei Isolation for Single Cell Multiome ATAC + Gene Expression Sequencing). Following nuclear isolation, GEMs were generated and barcoded using the Chromium Next GEM Chip J. ATAC and gene expression libraries were then constructed according to protocol CG000338, Revision E by 10X Genomics (Chromium Next GEM Single Cell Multiome ATAC + Gene Expression). Library size was subsequently quantified using a BioAnalyzer. All samples were sequenced on the Illumina NovaSeq 6000 system, with a target of 5 x 10^4^ RNA reads/nucleus and 2.5 x 10^4^ ATAC reads/nucleus. Sequencing reads were aligned to the human reference genome and pre-processed using the CellRanger v2.0.0 pipeline.

Downstream analysis of scMultiome data was conducted using the Seurat and Signac packages in R (Hao et al., 2021; Stuart et al., 2021). Low quality cells were first filtered out according to metrics reported in supplementary table 2A. Next, peaks were called using MACS2. Gene expression data were then normalized using SCTransform, with differences between timepoints being regressed out. Doublets were identified using the DoubletFinder package with the following parameters (pK = 0.1, pN = 0.05 (day 0), 0.01 (day 7), 0.005 (days 14 and 56), nExp_poi = 0.075 * number of cells in object), and then removed. Dimensionality was reduced with principal component analysis (PCA). DNA accessibility data were processed by latent semantic indexing (LSI). A multimodal nearest neighbour graph was generating using the top 50 PCs and top 40 LSI, after which a joint UMAP plot was constructed to visualize both data modalities. Individuals timepoints were clustered at a resolution of 0.2, and DE analysis was conducted between clusters using a log2 fold change of 0, and a maximum of 1000 cells per identity using a Wilcoxon Rank Sum test, and multiple hypothesis testing was corrected for. DE was also conducted between timepoints using the same parameters. Cell clusters were annotated using markers identified through DE (see supplementary table 2B). Differential accessibility (DA) analysis was performed between day 0 and days 7, 14 and 56 using above parameters. DA regions in each differentiation timepoint relative to day 0 with p < 0.005 were used to search for transcription factor motifs present in these regions. Motif activity scores were then computed using chromVAR (Schep et al., 2017).

### Statistical Methods

Data were analyzed using GraphPad Prism 9. Two-tailed parametric t-tests were performed to compute p values, and a threshold of p < 0.05 was used for statistical significance (*p < 0.05, **p < 0.01, ***p < 0.001). Data are represented as mean ± SEM, unless otherwise specified.

### Ethics Statement

All experiments with human cells were approved by The Hospital for Sick Children Research Ethics Board (REB# 0020020272).

## Acknowledgements

Research was supported by a Canadian Institutes of Health Research Project Grant (462770) and b.r.a.i.n.child grant (BC-19-10) awarded to P.B.D. B.M.D. was supported by a Frederick Banting and Charles Best Canada Graduate Scholarships – Doctoral award from CIHR. F.M. was supported by an Autism Speaks Award. J.E. holds a Tier 1 Canada Research Chair in Stem Cell Models of Childhood Disease and acknowledges support from the Canada Foundation for Innovation John R. Evans Leadership Fund. X. H. holds a Tier 2 Canada Research Chair in Cancer Biophysics. P.B.D. holds a Harold Hoffman/Shoppers Drug Mart Chair in Paediatric Neurosurgery and is also supported by the Hospital for Sick Children Foundation, Jessica’s Footprint, Hopeful Minds, and the Bresler Family.

---

**Supplementary figure 1.**
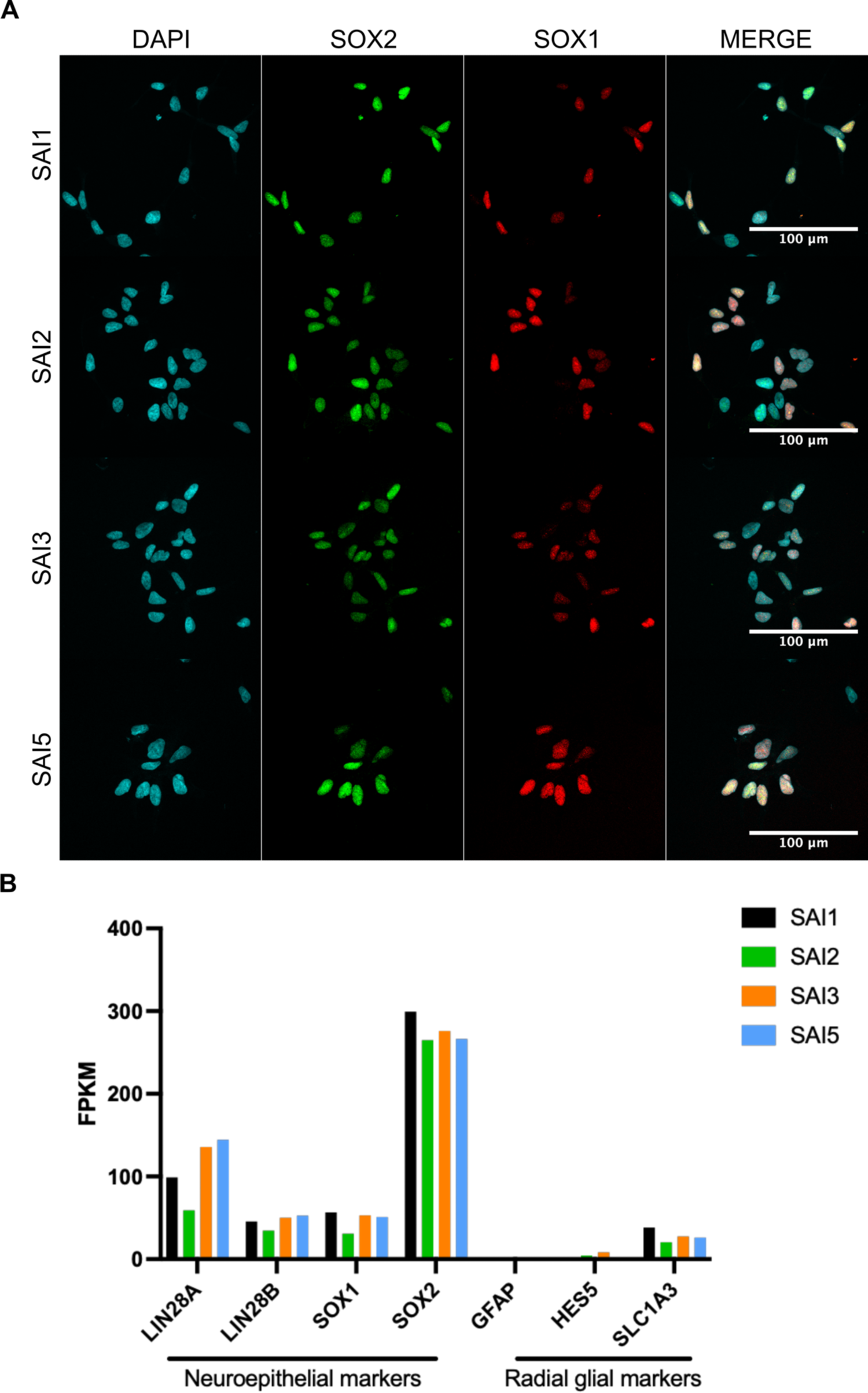
Undifferentiated NES cells stably retain regional and temporal identity *in vitro*. **A** Representative ICC images of neuroepithelial transcription factor SOX1 and SOX2 expression in undifferentiated NES cells. **B** Expression of neuroepithelial and glial markers in NES cells measured by bulk RNA-seq. Scale bars represent 100 μm.

**Supplementary Figure 2.**
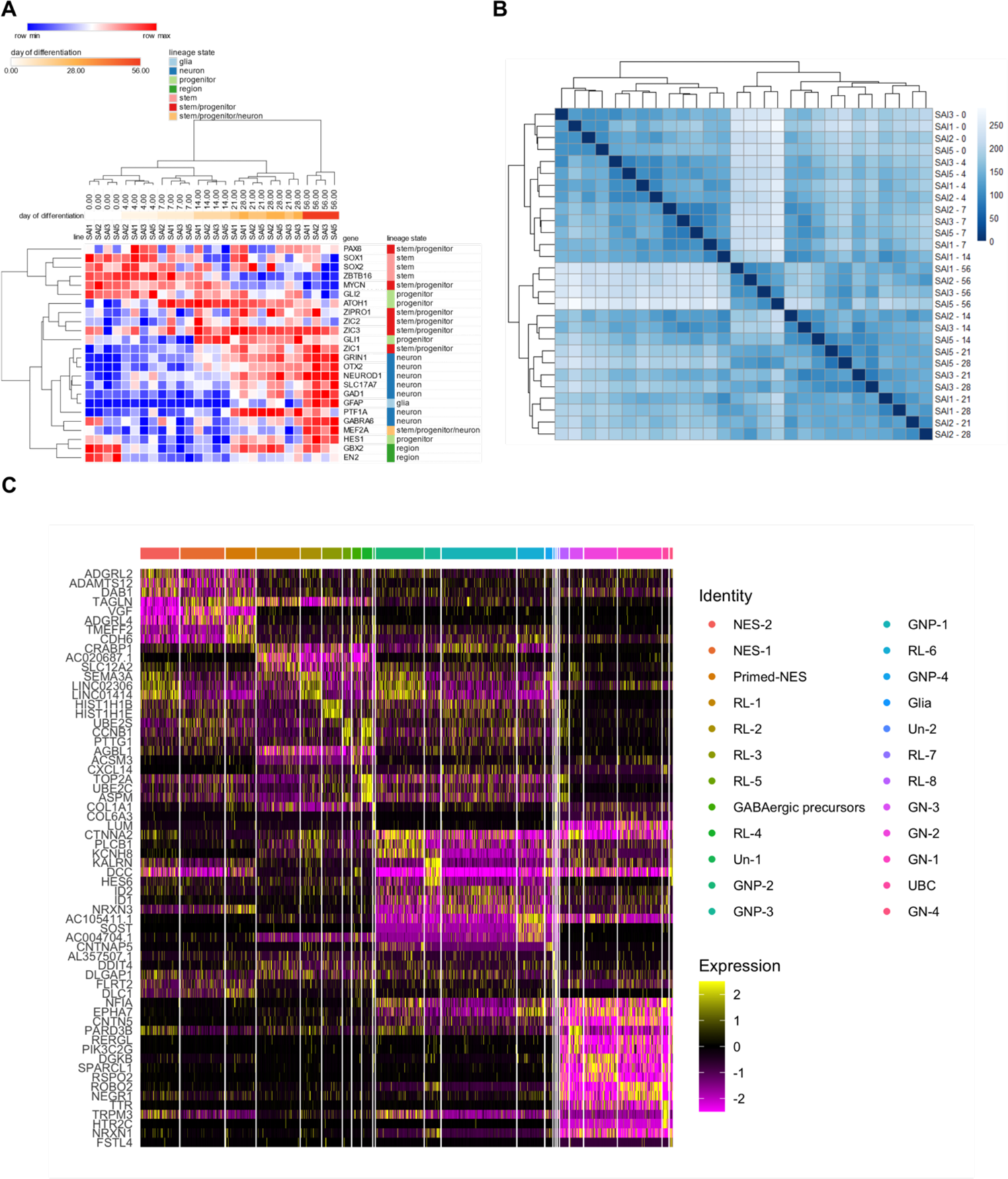
Directed differentiation of NES cells to cerebellar granule neurons. **A** Hierarchically clustered relative expression values of select granule lineage markers as measured by qRT-PCR. **B** Euclidean distance matrix of bulk transcriptomes during differentiation. **C** Heatmap of differentially expressed genes between all clusters annotated in figure 1F (see also supplementary table 2H).

**Supplementary figure 3.**
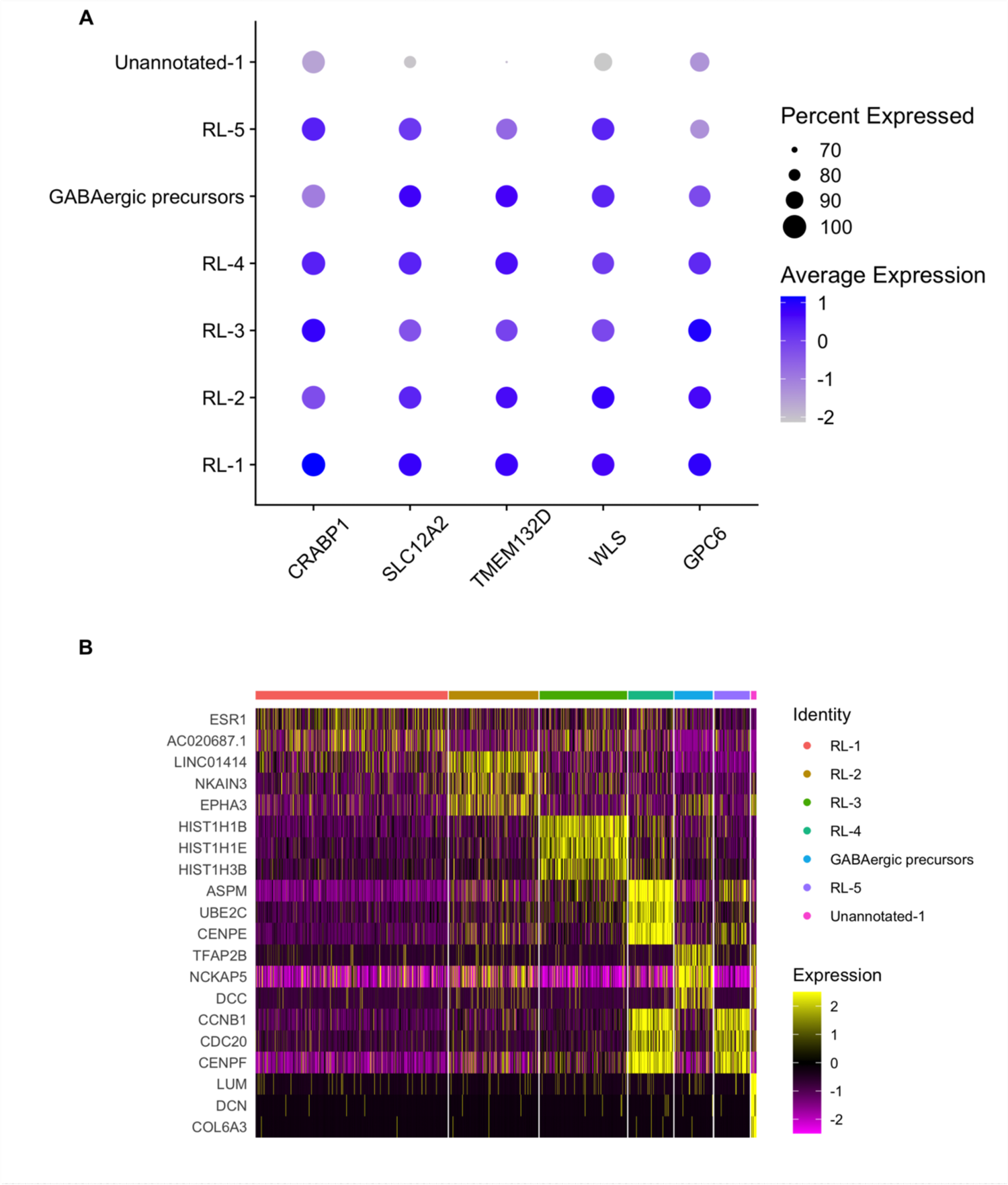
Emergence of a rhombic lip-like state at day 7 of differentiation. **A** Expression of rhombic lip markers among day 7 cell clusters. **B** Differentially expressed genes among day 7 clusters.

**Supplementary figure 4.**
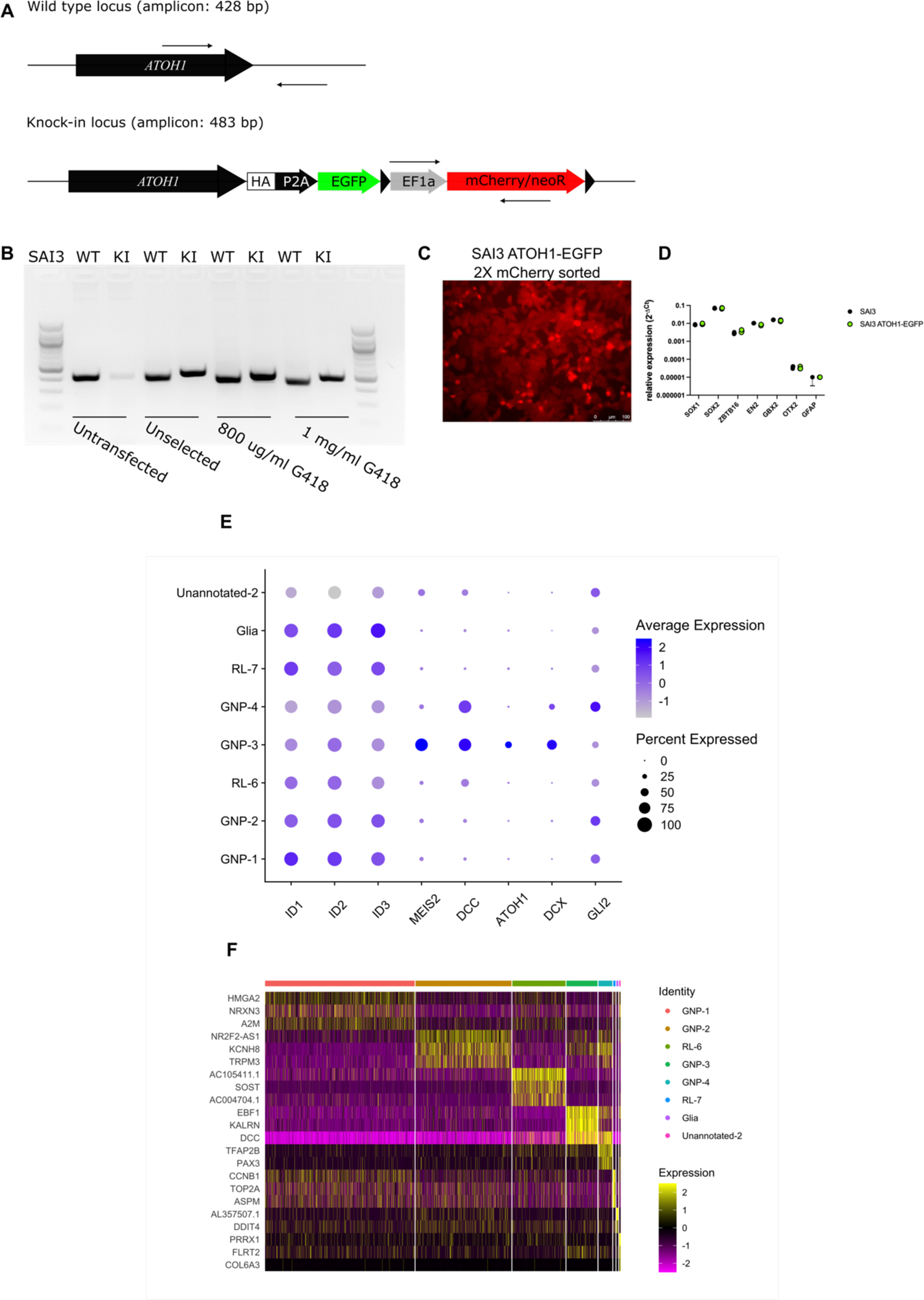
Design of an *ATOH1*-EGFP knock-in reporter and further characterization of day 14 cell clusters. **A** Design of homology repair template to introduce knock-in. Arrows indicate approximate positions of PCR primers (see Materials and Methods for primer sequences). **B** Genotyping PCR following nucleofection of gRNA/Cas9 plasmid and repair template and G418 selection. **C** Isolation of knock-in cells by twice sorting for mCherry^+^/DAPI^-^ cells. **D** Expression of key NES markers by qRT-PCR in parental SAI3 and SAI3 ATOH1-EGFP cells (n = 3 biological replicates). **E** Expression of select GNP markers among day 14 cell clusters (see also supplementary table 2J). **F** Differential gene expression analysis between day 14 cell clusters (see also supplementary table 2E).

**Supplementary figure 5.**
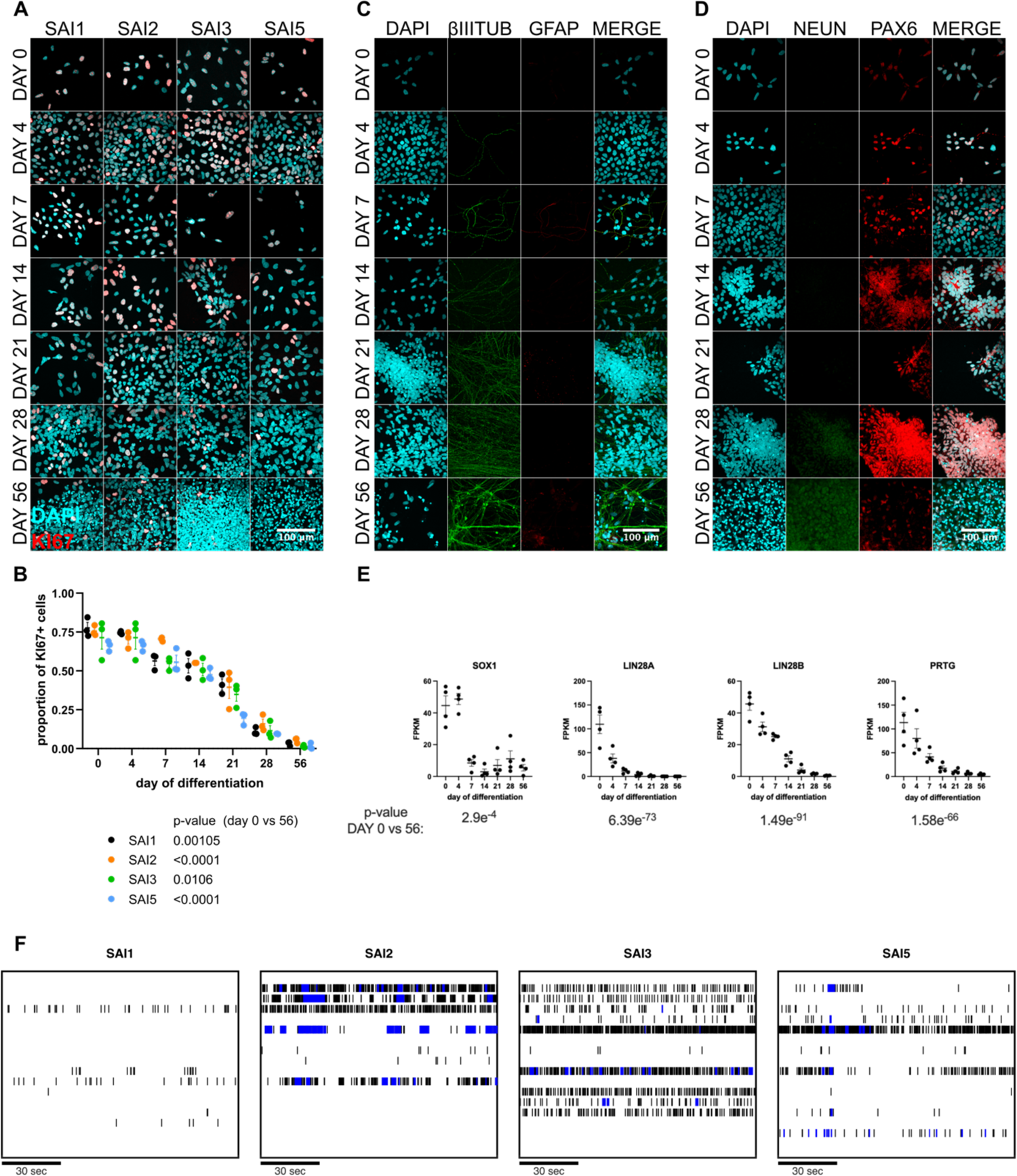
Neuronal arborization and loss of proliferation and stemness accompany granule neuron differentiation. **A** Representative ICC images of KI67 expression at various differentiation timepoints. **B** Quantification of proportion of KI67^+^ cells in A (n = 3 biological replicates, two-sided t-test). **C** Representative ICC images of neural (*β*III tubulin) and glial (GFAP) marker expression during differentiation in the SAI5 NES cell line. **D** Representative ICC images of NEUN (post-mitotic neurons) and PAX6 (granule lineage) expression during differentiation in the SAI5 NES cell line. **E** Bulk RNA-seq expression of neuroepithelial genes during differentiation. All scale bars represent 100 μm. **F** Representative raster plots of day 56 neuronal activity. Bursts are highlighted in blue. Scale bars represent 30 seconds.

**Supplementary figure 6.**
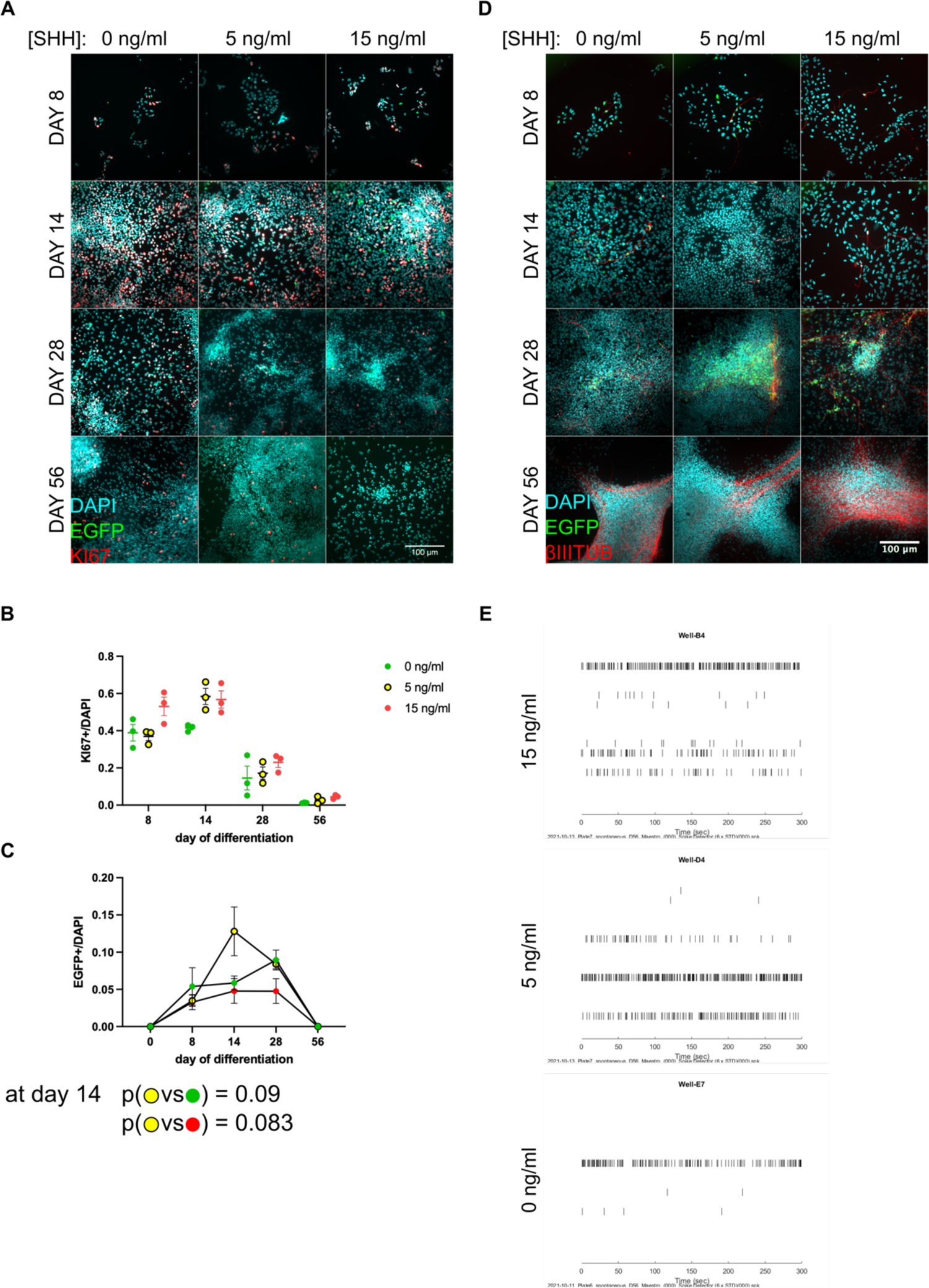
Testing the effects of SHH on granule neuron differentiation. **A** Representative ICC images of EGFP and KI67 expression during differentiation in varying SHH concentrations. **B** Quantification of KI67^+^ cell proportion in A. **C** Quantification of EGFP^+^ cell proportion in A. **D** Representative ICC images of EGFP and *β*III tubulin during differentiation. N = 3 biological replicates for all experiments, t-tests conducted for statistical significance. Scale bars represent 100 μm. **E** Representative raster plots of neuronal activity under various SHH concentrations.

